# Structure-guided inhibition of the cancer DNA-mutating enzyme APOBEC3A

**DOI:** 10.1101/2023.02.17.528918

**Authors:** Stefan Harjes, Harikrishnan M. Kurup, Amanda E. Rieffer, Maitsetseg Bayarjargal, Jana Filitcheva, Yongdong Su, Tracy K. Hale, Vyacheslav V. Filichev, Elena Harjes, Reuben S. Harris, Geoffrey B. Jameson

## Abstract

The normally antiviral enzyme APOBEC3A^1-4^ is an endogenous mutagen in many different human cancers^5-7^, where it becomes hijacked to fuel tumor evolvability. APOBEC3A’s single-stranded DNA C-to-U editing activity^1, 8^ results in multiple mutagenic outcomes including signature single-base substitution mutations (isolated and clustered), DNA breakage, and larger-scale chromosomal aberrations^5-7^. Transgenic expression in mice demonstrates its tumorigenic potential^9^. APOBEC3A inhibitors may therefore comprise a novel class of anticancer agents that work by blocking mutagenesis, preventing tumor evolvability, and lessening detrimental outcomes such as drug resistance and metastasis. Here we reveal the structural basis of competitive inhibition of wildtype APOBEC3A by hairpin DNA bearing 2’-deoxy-5-fluorozebularine in place of the cytidine in the TC recognition motif that is part of a three-nucleotide loop. The nuclease-resistant phosphorothioated derivatives of these inhibitors maintain nanomolar *in vitro* potency against APOBEC3A, localize to the cell nucleus, and block APOBEC3A activity in human cells. These results combine to suggest roles for these inhibitors to study A3A activity in living cells, potentially as conjuvants, leading toward next-generation, combinatorial anti-mutator and anti-cancer therapies.

## INTRODUCTION

The APOBEC3 family of enzymes (A3A-A3H) forms part of the innate immune response against viruses and transposons^1-4^. These enzymes deaminate cytosine to uracil in single-stranded DNA^1, 8^ (ssDNA) and, a subset, also in RNA^10-12^. In particular, A3A and A3B promote mutagenesis of genomic DNA fueling cancer evolution leading to drug resistance and poor disease outcomes^5-7^. Recent studies show that A3A is capable of driving carcinogenesis in mice^9^ and contributing the majority of APOBEC signature mutations to human cancer cells^12-13^. Therapeutic applications of A3 inhibitors have been proposed as a strategy to prevent A3-mediated evolution of primary tumors into lethal metastatic and drug-resisting secondary growths^14-15^.

A3A preferentially deaminates the 5*′*-YT<u>C</u>R target motifs, whereas A3B targets 5’-NT<u>C</u>R, where N denotes A, C, G, or T, Y pyrimidine, and R purine^12-14, 16-17^. The preference of these enzymes for the T<u>C</u> dinucleotide recognition motif is explained by previous crystal structures of linear ssDNA bound to an inactive mutant of A3A, which show binding pockets for the target cytosine and preceding thymine^18-22^. However, this leaves the preference for the nucleotides flanking the T<u>C</u>-sequence unexplained. Crystallographic and NMR studies have revealed that upon binding to A3A the ssDNA adopts a distinctive *U*-shape that projects cytosine into the A3A active site^18-19, 23^. Consistent with this observation, DNA hairpins with loops of three and four nucleotides have been shown to be more potent substrates of A3A in comparison to the linear analogs^24-28^. Importantly, DNA hairpins are both physiologically and pathologically relevant. Such structures are ubiquitous in nature, especially at inverted repeats, which cause stalling of replication, genomic instability, and predisposition to mutagenesis^28-29^.

Prior studies have shown that linear ssDNA substrates with 2*′*-deoxyzebularine (dZ) and 5-fluoro-dZ (FdZ) in place of the target cytosine are weak inhibitors of A3A (and A3B)^30-32^. Additional work by our group and others has shown greater inhibition of A3A *in vitro* with pre-formed *U*-shaped, hairpin dZ and FdZ transition-state trapping oligonucleotides^33-34^. Here, we use X-ray crystallography to reveal the first high-resolution structures of wildtype A3A-inhibited complexes and demonstrate the underlying mechanism of inhibition. Importantly, these structures demonstrate why 3-nucleotide hairpin loops with TTC (or TTFdZ) are preferred substrates (or inhibitors) for A3A. Larger 4-nucleotide loops extrude one nucleobase to optimally fit into the A3A active site. Moreover, here we also demonstrate for the first time that a FdZ hairpin, TTFdZ-hairpin, is not only a potent nanomolar inhibitor *in vitro* in biochemical assays but also a potent inhibitor of wildtype A3A-catalyzed chromosomal DNA editing activity in human cancer cells.

## RESULTS

### FdZ is hydrated in the active site of A3A and mimics an intermediate of cytosine deamination

The mechanism for CDA deamination was proposed based on several crystal structures in which zebularine and 5-fluorozebularine accept a Zn^2+^-bound water molecule across the N3-C4 double bond forming a tetrahedral intermediate when complexed with CDA (**Fig. 1a**)^35-37^. To address whether A3A utilizes the same deamination mechanism, we crystallized wildtype A3A complexed with an inhibitory hairpin (TTFdZ-hairpin: 5*′*-TGCGCTTFdZGCGCT; **Fig. 1b**). Two slightly different crystal forms were obtained and resolved to 2.80 and 2.94 Å (**Extended Data Fig. 1a**). **Fig. 1c** shows that the N3-C4 double bond of FdZ is hydrolyzed so that the hydroxy group at C4 represents the tetrahedral intermediate in the deamination of cytosine – here with defined *R*-stereochemistry, 4-(*R*)-hydroxy-3,4-dihydro-2’-deoxy-5-fluorozebularine. The ^−^OH at C4 is bound to the Zn^2+^ center and is derived from the water/hydroxide ion bound to the Zn^2+^ in the substrate-free state. Glu72, which functions as a general acid/base, hydrogen bonds (strictly forms a semi-salt bridge) to C4-OH and N3-H of hydrated FdZ. We expect for such interactions that Glu72 is present in the carboxylate form in the crystal structure. The 5-fluoro group of FdZ is accommodated comfortably, abutting Tyr130 (similarly to cytosine depicted in **Fig. 1g**). This, together with prior work indicating hydration of dZ and its derivatives in structures with CDA, A3A^19-20^ and A3G^38-40^, demonstrate a universal mechanism of inhibition^18-19, 38-40^.

**Fig. 1:**
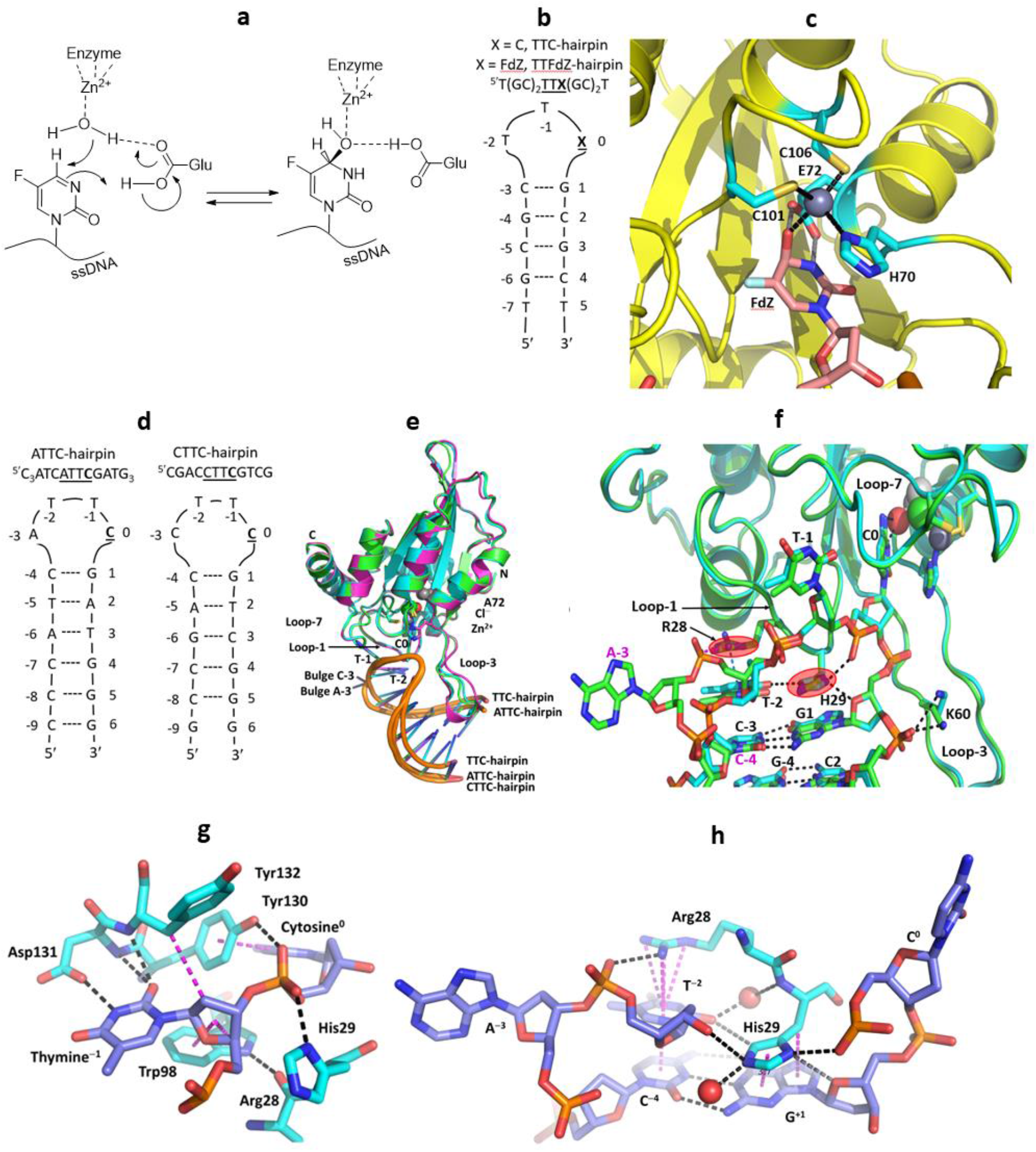
Structures of substrates and inhibitor of A3A, and mechanism of inhibition. **a**, Schematic of reaction of FdZ with water activated by Zn^2+^, analogous to that of cytidine/cytosine deaminases, showing critical role of general acid-base Glu72. **b**, TTFdZ- and TTC-hairpins. **c**, Structure of wildtype A3A with TTFdZ-hairpin inhibitor with FdZ hydrolyzed and coordinated to the active-site Zn^2+^. Glu72 and metal-binding ligands (His70, Cys106 and Cys106) are highlighted in cyan. **d**, ATTC- and CTTC-hairpin substrates of A3A with four-residue loops. **e**, Superposition of structures of A3A-E72A in complex with three-residue loop TTC-hairpin (cyan) and four-residue loop ATTC-hairpin (green) and CTTC-hairpin (magenta). **f**, Superposition of A3A-E72A in complex with TTC-hairpin (cyan) and four-residue loop ATTC-hairpin (green). The critical roles played by His29 and Arg28 in determining the conformation of the loop that makes these hairpins better substrates for wildtype A3A than corresponding linear ssDNA are highlighted. **g**, Interactions of Loop-7 and Loop-1 residues of A3A-E72A with the TC recognition motif. **h**, Interactions of Arg28 and His29 of Loop 1 of A3A-E72A with the ATTC-hairpin. Key hydrogen-bonding interactions are shown by black dashes. Cation-*π* interactions between Arg28 and T^−2^, *π*-*π* interactions between T^−2^ and G^−4^ and between His29 and G^+1^, and dispersion interactions are shown as magenta dashes. Zn^2+^ are shown as grey spheres and Cl^−^ as green spheres. All structures share a similar crystal packing, whereby the base of the stem of one molecule stacks with that of the other molecule in the asymmetric unit (**Extended Data Fig. S2**). (246/250 words)

To ascertain generality of the interactions observed between hairpin DNA and A3A, we determined structures of A3A as its inactive E72A mutant in complex with three distinct hairpins (**Fig. 1b,d**). All structures share a similar crystal packing (**Extended Data Fig. 2**), a common tertiary structure for the protein, and a common conformation of the hairpins’ TT(C/FdZ)G moiety (**Extended Data Fig. 3**). The superpositions of A3A-E72A in complexes with TTC-hairpin (at 2.22 Å resolution) and the four-residue loop ATTC-hairpin (at 1.91 Å resolution) and CTTC-hairpin (at 3.15 Å resolution) are shown in **Fig. 1e. Fig. 1f** shows the top of the hairpin stem and along with the TTC and ATTC loops their conserved interactions with residues Arg28 and His29. The structures of wildtype A3A (two near-isoforms) and A3A-E72A, along with their respective hairpin DNA, all superimpose very closely (average RMSD of 0.43 Å), noting, in particular, the very close superposition of the TTC moiety with the TTFdZ moiety (**Extended Data Fig. 1**). The only exception to close superposition of the hairpins with three- and four-nucleotide loops is that the residue at position −3 of the four-residue loop is flipped out, but the next residue C^−4^ is in register to hydrogen bond with dG^+1^, as in structures with a three-residue loop. Further details of these and other structures are provided in **Extended Data** (Fig. 1-8 and Tables 1 and 2) and in **Supplementary Information**, where additional analysis and discussion of results may be found.

### His29 and Arg28 play critical roles determining potency of hairpin substrates and TTFdZ hairpin inhibitor toward A3A

In contrast to existing structures of linear ssDNA with A3A mutants^18-19^, our structures reveal the basis for higher activity of hairpin DNA substrates and the expected greater potency of corresponding inhibitors. First, His29 base-stacks with G^+1^ and T^−2^ base-stacks with the pyrimidine in the stem (C^−3^ for the three-residue loop or for the four-residue loops C^−4^) (**Fig. 1g,h**, and **Extended Data Fig. 3a-c**). Second, the tight turn between the nucleotide at position +1 and the target cytosine at position 0 is stabilized by a bifurcated hydrogen bond between Nδ_1_ (amine tautomer) of His29 and O4 of 2-deoxyribose at +1 and an oxygen atom of the phosphate group that links nucleotides C^0^ and T^−1^. Third, a non-classical hydrogen bond exists between atom Cd_2_ of His29 and the O2 atom of C=O in T^−2^ (distances 3.2-3.4 Å) further stabilizing the non-canonical His-thymidine pair. This provides an additional base-pair in which one base is from DNA and another is from the protein, both stacking on top of the first CG base pair (at positions −3 or −4 and +1) at the head of the stem. This first CG base-pair shows a small distortion from coplanarity. By virtue of chemical structures, purines are larger than thymine and have nothing to resemble the C=O of T or C to form the crucial H-bond with His29. Therefore, this non-canonical base-pairing explains why A3A prefers pyrimidines in the –2 position, as reflected in whole genome sequences of A3A-mutagenized model systems.^9, 13, 17^ Last, a water molecule bridges the carbonyl O2 atom of T^−2^ and peptide N of His29 (**Fig. 1h**).

With regard to the role of Arg28, its positively charged guanidinium moiety forms a cation-*π* interaction with the nucleobase T^−2^ thus further stabilizing T^−2^ placement for non-canonical base-pairing with His29. When the loop length is increased from three to four nucleotides, the additional nucleotide in position −3 is flipped out preserving vital His29/T^−2^ interactions in these hairpins. In these structures Arg28 forms a hydrogen bond to an oxygen atom of the phosphate linking A^−3^/C^−3^ to T^−2^ (**Fig. 1h**), and the cytosine at position −4 is in register to hydrogen bond with guanine at position +1 as a part of hairpin’s stem.

A3 enzymes prefer DNA over RNA structures and, in accordance, all deoxyribose moieties adopt the standard for DNA C2-*endo* conformation in our structures, with possible exceptions for nucleobases at the 5’ and 3’ ends of the stem, which are poorly defined in electron density maps. The tight turn to project C^0^ into the active site is accomplished with non-standard torsional angles for the phosphate groups, relative to expected values for A- or B-DNA, as detailed in **Supplementary Information**.

### TTC-hairpin is a better substrate for wildtype human A3A than linear ssDNA

Hairpins with the target cytosines at 3’ end of the three-nucleotides TTC-loop are the preferred substrates for A3A, as longer loops are deaminated with lower efficiency^25^. For kinetic characterization we have chosen the hairpin with the shortest stem of four base-pairs showing maximal deamination. A CG base-pair closing the loop is also preferred in the stem^25^. To make sure that DNA hairpin is thermodynamically stable we used three further CG/GC base-pairs in the stem (**Fig. 1b,d**). CD and NMR spectra confirm that a hairpin structure forms in solution (**Extended Data Fig. 3d,e**).

Our real-time NMR assay^41^ compared deamination of TTC-hairpin and linear ssDNA by wildtype A3A in a buffer at pH 7.4 over the complete time course of product formation (**Fig. 2a**). This enabled use of the integrated Michaelis-Menten equation by means of Lambert’s W function, which provides more robust estimates for kinetic parameters *K*_m_ and *V*_max_ than analysis of initial rates only.^42^ Kinetic constants obtained (**Fig. 2b**, see also Supplementary Information) indicate that TTC-hairpin is a preferred substrate of A3A in our experimental conditions, which agrees with previously reported data of a fluorescently-based assay^13, 25, 28^. A dramatic change in *K*_m_ from 3.0 ±0.9 mM for linear ssDNA to 31 ± 6 *μ*M for TTC-hairpin is the major contributor to a 42 times more efficient deamination of TTC-hairpin if *k*_cat_ / *K*_m_ are compared at pH 7.4.

**Fig. 2:**
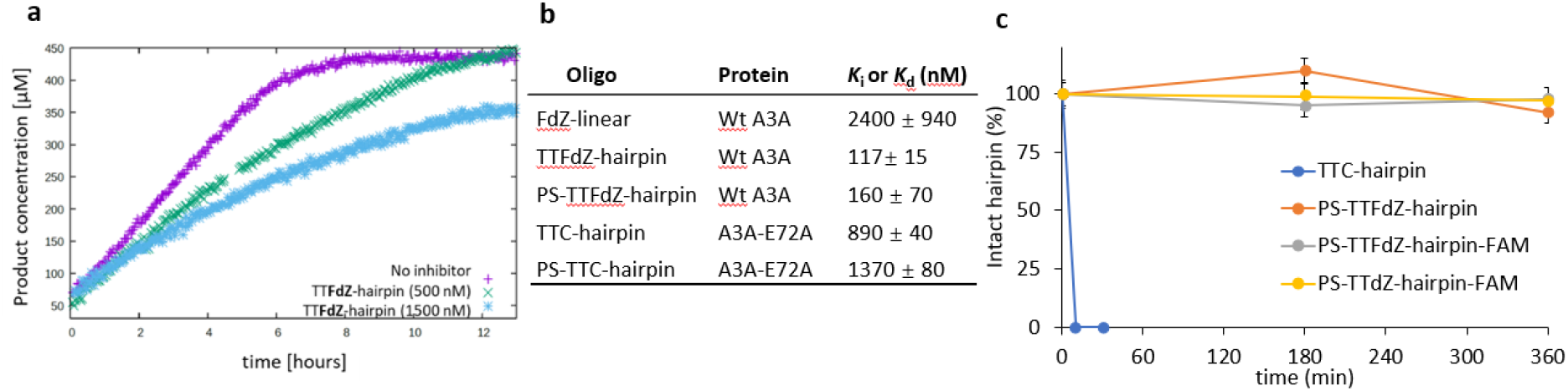
Kinetics of A3A-catalyzed deamination of DNA-hairpin and susceptibility of hairpin DNA to degradation by snake venom phosphodiesterase. **a**, Plot of product concentration in the absence and presence of TTFdZ-hairpin at 500 and 1500 nM; concentration of A3A 140 nM; concentration of TTC-hairpin substrate 500 *μ*M. **b**, Table (upper 3 entries) shows derived inhibition constants, *K*_i_ for linear FdZ ssDNA (A_2_T_2_FdZA_4_), TTFdZ-hairpin and its phosphorothioated analog PS-TTFdZ-hairpin. Table (lower 2 entries) shows *K*_d_ obtained by isothermal titration calorimetry (ITC) for binding of TTC-hairpin and PS-TTC-hairpin to A3A-E72A. **c**, Percentage of intact hairpins (15 μM) over time upon treatment with snake venom phosphodiesterase (phosphodiesterase I, Sigma, 32 mU/mL) in 50 mM Tris-HCl buffer, 10 mM MgCl_2_, pH 8.0 for the indicated times at 37 °C. Full experimental details are available in Supporting Information.

### TTFdZ-hairpin is a better A3A inhibitor than linear ssDNA

The exchange of the target C with a nucleoside-like CDA inhibitor FdZ (**Fig. 1a,b**) in a DNA hairpin led to a potent inhibition of wildtype A3A in comparison with the linear FdZ-oligo^25, 30-32^ (**Fig. 2a,b**). The obtained *K*_i_ values (**Fig. 2b** and **Extended Data Fig. 4a**) show that TTFdZ-hairpin is a superior inhibitor of wildtype A3A in comparison with the best FdZ linear ssDNA previously described^31^. These data support our hypothesis that better substrates can be converted into the better inhibitors. Taken together, the interactions between His29 and Arg28 with DNA become more defined when hairpins are used explaining the YTCR-recognition motif and the superior properties of hairpin DNA as substrate and, appropriately modified, as inhibitor compared to corresponding linear ssDNA species.

### Phosphorothioated (PS) hairpins inhibit A3A *in vitro* and are resistant to nucleases

For testing in cells, oligonucleotides were prepared with thiophosphate (PS-) where all (normal) O-P(O)_2_-O linkages replaced with O-P(OS)-O (denoted PS-TTC-hairpin and PS-TTFdZ-hairpin). This modification, commonly used in oligo therapeutics^43^, protects DNA against nucleolytic degradation that can occur within minutes in cells. To test that the PS modification does not alter the binding to A3A, isothermal titration calorimetry (ITC, see Supporting Information) was performed with the TTC-hairpin and PS-TTC-hairpin and an inactive mutant of A3A, A3A-E72A, where the active-site glutamic acid is replaced with alanine to prevent unwanted deamination of the substrate. The PS-TTC-hairpin binds to A3A-E72A with only a slight decrease in affinity compared with that for TTC-hairpin with native PO linkages (**Fig. 2b, Extended Data Fig. 4a,b**), despite many diastereomeric forms (2^12^) present in PS-TTC-hairpin. In accord with these results, FdZ-containing hairpin with PS linkages (PS-TTFdZ-hairpin) inhibited wildtype A3A similarly to the TTFdZ-hairpin with PO-linkages (**Fig. 2b**). To demonstrate biostability of these PS-modified inhibitors, we analyzed the resistance of the PO and PS-hairpins to snake venom phosphodiesterase, an enzyme with strong 3’-exonuclease activity that is commonly used to evaluate the stability of oligos having therapeutic potential^44^. As shown in **Fig. 2c**, whereas the TTC-hairpin was completely degraded within 10 min, the PS-containing hairpins demonstrated enhanced stability against enzymatic digestion, remaining fully intact after 6 hours.

### Phosphorothioated (PS-) hairpin localizes to the cell nucleus and inhibits A3A activity *in cellulo*

In cell lysate, PS-TTFdZ-hairpin inhibits, in a concentration-dependent manner, the activity of A3A on TTC-hairpin substrate (**Fig. 3a,b**). The PS-TTT-hairpin as control showed no inhibition of A3A activity. To assess subcellular localization, the modified PS-hairpins possessing dZ or FdZ were fluorescently labelled at the 3’-end (6-FAM). These FAM-labelled PS-hairpins are also resistant to 3*′*-exonuclease activity (**Fig. 2c**). MCF-7 breast cancer cells were transfected with these hairpins using the Xtreme GENE™ HP transfection agent. After 18 hours these FAM-labelled PS-hairpins localize to the nucleus in a concentration-dependent manner (**Fig. 3c** and **Extended Data Fig. 5a-e**). Cell viability was tested on two breast cancer cell lines (MCF-7 and MDA-MB-453) and found to be dependent on hairpin concentration, decreasing to 50-70% viable at 20 *μ*M concentration of PS-TTT-hairpin or PS-TTFdZ-hairpin. On the other hand, cell viability was unchanged between 24 and 48 h (**Extended Data Fig. 6**). These data demonstrate the benefit of introducing the PS-linkage, as it results in hairpins that are stable in biological media while maintaining efficiency of A3A inhibition.

**Fig. 3:**
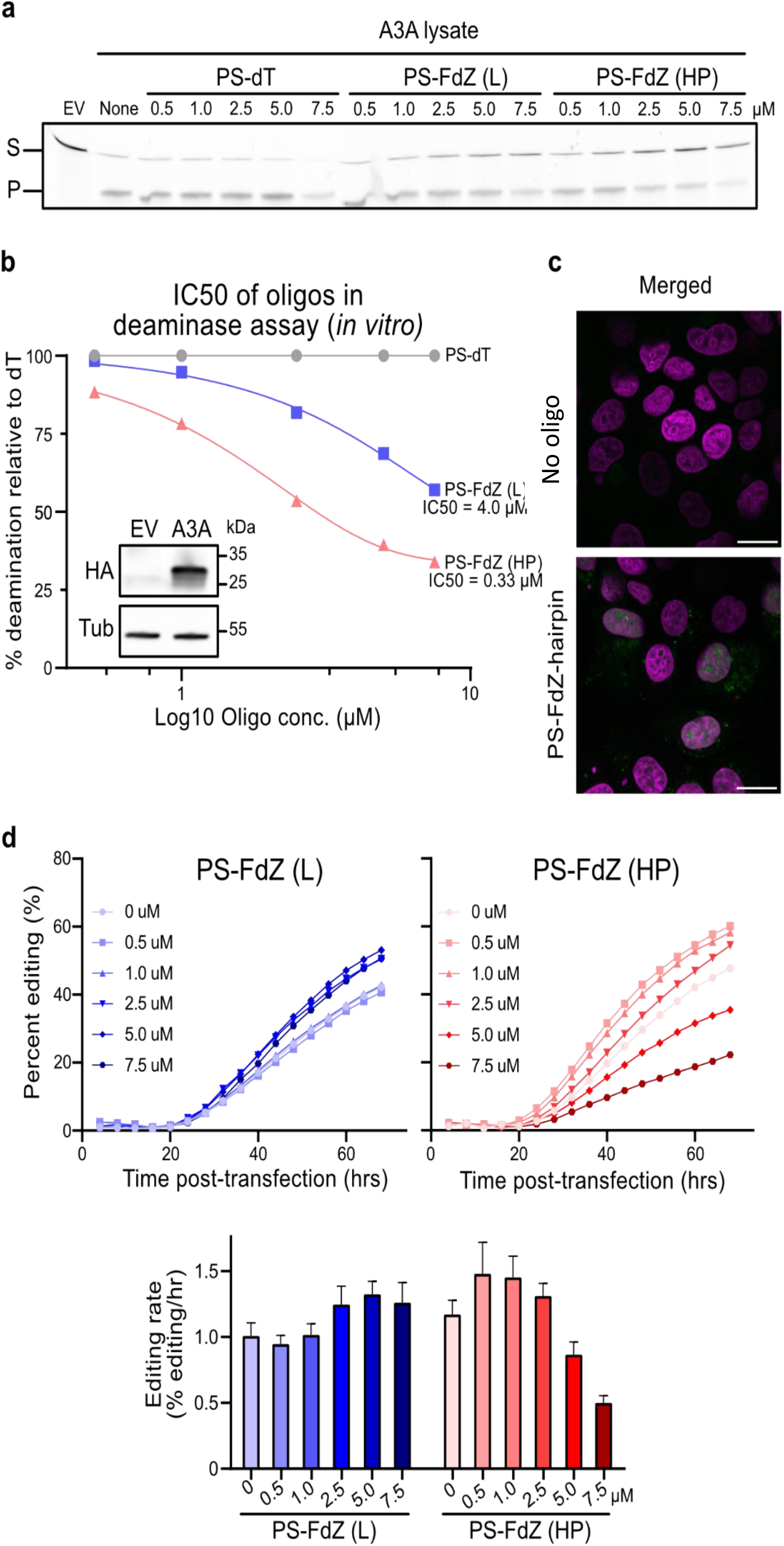
*In cellulo* properties of fluorescently tagged PS-TT(F)dZ-hairpin-FAM. **a**, Concentration dependence of A3A activity in cell lysates in presence of PS-TTT-hairpin (denoted PS-dT), PS-TTFdZ-linear (denoted PS-FdZ (L)), and PS-TTFdZ-hairpin (denoted PS-FdZ (HP)). **b**, Bands from panel (**a**) were quantified, normalized to the dT oligo, and used to determine the IC50 for FdZ (L) and FdZ (HP). The inset panel is an immunoblot showing expression of A3A-HA from cell lysate. A3A-HA was detected by an anti-HA antibody while anti-tubulin was used as a loading control. **c**, Representative images of asynchronously-grown MCF-7 cells transfected using Xtreme GENE™ HP with either no hairpin (top panel) or 1.25 μM PS-TTFdZ-hairpin (bottom panel) with PS-linkages. MCF-7 cells were incubated for 16 hours with hairpin and Xtreme GENE ™ HP. Images have the pseudo-coloured panels overlaid: nucleus (magenta) and oligo-FAM (green). Variability in hairpin uptake is attributed to cells being at different stages of their cycle. Scale bars, 20 *μ*m. **d**, PS-TTFdZ-hairpin shows concentration-dependent inhibition of A3A-editing activity, whereas weakly binding linear PS-FdZ (L) and non-A3A-binding PS-TTT-hairpin have negligible effects on evolution of GFP fluorescence (**Extended Data Fig. 8**). Biological replicates, establishing reproducibility, are shown in **Extended Data Fig. S8**. Variability in plasmid transfection prevents averaging of results. The transfection reaction (TransIT-LT1), administered at constant concentrations across all *in cellulo* experiments in presence or absence of DNA species, appears to have a slight activating effect on A3A editing activity at low concentrations of both hairpin and linear DNA. (242/∼250 words)

*In cellulo* inhibition of A3A activity was quantified using a non-covalent variation of a base editing system reported previously^45-46^. Briefly, A3A-catalyzed editing of a single target cytosine nucleobase within a ssDNA R-loop created by a nuclease-deficient Cas9-guide RNA complex in a *eGFP* reporter construct results in a dose- and time-dependent restoration of eGFP fluorescence (**Methods** and **Extended Data Fig. 7**). This reporter has been stably integrated into 293T cells to better mimic untethered A3A mutagenic deamination at the chromosomal level. Upstream of the mutated *eGFP* codon lies a linked wildtype *mCherry* gene for calculating the efficiency of base editing (eGFP+/mCherry+). With PS-TTT-hairpin as a A3A-non-binding control, little change in generation of GFP fluorescence with time is observed as a function of concentration (**Extended Data Fig. 8a,b**). However, in the presence of increasing concentrations of PS-TTFdZ-hairpin inhibitor, there is marked suppression of the generation of fluorescence (**Fig. 3c**). The maximum rate of A3A inhibition is 2.4-fold at 7.5 *μ*M concentration of inhibitor compared to absence of inhibitor, suggesting an upper limit on IC_50_ of ∼5 *μ*M. The difference in cell viability between PS-TTT and PS-TTFdZ hairpins is minimal in comparison with A3A inhibition data for these oligos at the same concentrations in 293T cells (**Extended Data Fig. 8** and **Fig. 3c**).

## CONCLUSIONS AND PERSPECTIVES

Our structural studies establish that the stem-loop preconfigures the TC recognition motif in optimal position for binding to A3A, such that hairpin DNAs are more active substrates and as the 2*′*-deoxy-5-fluorozebularine derivative more potent inhibitor of A3A than linear ssDNA. Both 3- and 4-membered nucleotide loops present the TC motif in identical configuration. Although hinted at in earlier structures with ssDNA, a crucial role for His29 (and to a lesser extent Arg28) in substrate binding is confirmed, and importantly here also uniquely explaining A3A preferential deamination of YTCR motifs^17^. This has implications for activity of A3A leading to genomic instability and for the design of inhibitors to mitigate mutagenic activity of A3A in a wide variety of cancer types. Interestingly, A3B does not have a strong preference in –2 position^13^ and has an Arg instead of His29 in the corresponding position^17^. As phosphorothioated derivatives, our inhibitors are highly resistant to nuclease degradation and can be directed to the nucleus with the aid of commonly used transfection reagents. Moreover, we have obtained the first proof of inhibition of the mutagenic activity of A3A in cells by a relatively small molecule.

## Supporting information

Additonal analysis and discussion of results

## METHODS

### Oligonucleotide Synthesis and Purification

The general strategy for the synthesis of dZ and FdZ and their incorporation into DNA oligomers has been described by us elsewhere^33^. 3-[(Dimethylaminomethylidene)amino]-3*H*-1,2,4-dithiazole-3-thione (DDTT, Sulfurizing Reagent II from GlenResearch, USA) was used for sulfurization of oligos. The sulfurization step (2 - 4 min) was conducted before capping as a replacement of the standard oxidation step. FAM is located at 3’-end of the oligo and was obtained by synthesizing an oligo on a controlled pore glass loaded with 4-[6-[(2*S*,4*R*)-4-hydroxy-2-(DMT-*O*-methyl)pyrrolidin-1-yl]-6-oxohexyl]carbamoylfluorescein purchased from PrimeTech (Cat. number: 008a-500, Minsk, Belorussia). dZ and FdZ phosphoramidites were synthesized as previously described^31-33^.

The final detritylated **dZ**-containing oligos were cleaved from the solid support and deprotected at r.t. using conc. NH_4_OH overnight. **FdZ**-Containing oligos were deprotected on the solid support by a two-step procedure with 10% Et_2_NH in CH_3_CN for 5 min, followed by incubation of the support in ethylenediamine/toluene mixture (1/1, v/v) for 2 hrs at room temperature.^47^ The support was washed with toluene (3 × 1 mL), dried *in vacuo* and the deprotected **FdZ**-containing oligo was released in H_2_O (1 mL).

The deprotected oligos in solution were freeze-dried and dry pellets were dissolved in milli-Q water (1 mL) and purified and isolated by i) reverse-phase HPLC on 250/4.6 mm, 5 μm, 300 Å C18 column (Thermo Fisher Scientific) in a gradient of CH_3_CN (0→20% for 20 min, 1.3 mL/min) in 0.1 M TEAA buffer (pH 7.0) with a detection at 260 nm or ii) ion-exchange (IE) HPLC using TSKgel Super Q-5PW column from TSK in buffer A [25 mM Tris°HCl, 20% CH_3_CN, 10 mM NaClO_4_, pH 7.4] and buffer B [25 mM Tris°HCl, 20% CH_3_CN, 600 mM NaClO_4_, pH 7.4]. Gradients: 3.7 min 100% buffer A, convex curve gradient to 30% B in 11.1 min, linear gradient to 50% B in 18.5 min, concave gradient to 100% B in 7.4 min, keep 100% B for 7.4 min and then 100% A in 7.3 min. Flow rate: 0.8 mL/min with a detection at 260 nm.

Oligonucleotides were freeze-dried, pellets were dissolved in milli-Q water (1.5 mL) and desalted by reverse-phase HPLC on a 100/10 mm, 5 μm, 300 Å C18 column (Phenomenex) in a gradient of CH_3_CN (0→80% for 15 min, 5 mL/min) in milli-Q water with detection at 260 nm. Pure products were quantified by measuring absorbance at 260 nm, analyzed by ESI-MS and concentrated by freeze-drying (**Table S1**).

### Expression and purification of A3A constructs

A3A-E72A was expressed and purified as described.^23^ A3A-E72A was used for structural studies and ITC experiments with the substrate. Wildtype A3A, which was recombinantly expressed with the His_6_ tag at the C-terminal end in *E. coli* and purified as described^33^, was used for kinetic and structural studies with the inhibitor. The yield from 1-10 L expression was usually not enough to justify size-exclusion chromatography purification. The protein was used in high-salt buffer for kinetics. For crystallography the oligonucleotide ligand was added in a 1:2 molar ratio and the protein buffer was changed to low-salt buffer using 3 times reconcentration with centrifugal filtration having 10 kDa cut off.

### Co-crystallization of A3A constructs with hairpin substrates

A3A-E72A (1-4 mM in low salt buffer) was mixed with oligonucleotides (10 mM in TE buffer: 10 mM Tris/HCl pH 7.9, 1 mM EDTA)) in a 1:2 molar ratio and diluted to 0.75 mM using above low salt buffer. Dilution was done with protein buffer. The mixture was added to crystallization solution in a 1:1 ratio and the mixture was pipetted on siliconized glass disks and sealed on top of a reservoir of crystallization solution for hanging-drop crystallization at 12 °C. The crystallization solution had the following composition: 100 mM bicine at pH 6.6, 200 mM NaCl, 20 mM putrescine, 1 mM TCEP, 1 mM inositol hexaphosphate (phytic acid) and 45 % pentaerythritol propoxylate (5/4 PO/OH). The Zn^2+^-free crystals of A3A-E72A with ssDNA were crystallized using A3A-E72A that had been purified in the presence of 1 mM EDTA.

His_6_-tagged wildtype A3A was mixed with 2 × molar ratio of inhibitor oligonucleotide and the buffer was changed to low salt buffer by washing 3 times using centrifugal filtration with 10 kDa cut-off. Then the protein concentration was adjusted to 0.85 mM and crystallization proceeded as described for substrates with A3A-E72A.

### X-ray crystallography

Notwithstanding two distinct crystal habits (tiny flattened needles and thin plates), all structures are approximately isomorphous with space group *P*2_1_ (*Z ′* = 2) and unit cells of dimensions *a* ≈ 52 Å, *b* ≈ 57 Å, *c* ≈ 92 Å and *β* ≈ 105°. Data were processed on-site at the Australian Synchrotron using XDS.^48-49^ Each structure was solved independently by molecular replacement (MolRep^50^) using the A3A structure PDB ID 5keg^19, 51^ (space group *I*222) from which metal ions, ssDNA, chloride ions and waters had been stripped. After rigid body refinement with REFMAC5^52^ of the CCP4 suite,^53^ initial electron density maps, visualized with COOT,^54^ showed clearly the presence, or in one structure absence, of Zn^2+^, along with remarkably well-defined electron density for the ssDNA hairpin. Structure elucidation proceeded with rounds of building with COOT and refinement with REFMAC5. In all structures, active site Loop-3 was not well defined, as well as Loop-2 that is remote to the active site. In several structures, phytic acid (inositol hexaphosphate) was ill-defined with only three phosphate groups being well-defined. **Table S2** presents a summary of crystallographic data, data collection and structure refinement. **Figures S1-S9** illustrate crystal packing, molecular structures and superpositions.

### Circular Dichroism (CD) Spectroscopy of Hairpin DNA

CD spectra were recorded using a Chirascan CD spectrophotometer (150 W Xe arc) from Applied Photophysics with a Quantum Northwest TC125 temperature controller. CD spectra (average of at least 3 scans) were recorded between 200 and 350 nm with 1 nm intervals, 120 nm/min scan rate and 10 mm path length followed by subtraction of a background spectrum (buffer only). CD spectra of G4-DNAs were recorded at ∼10 µM DNA concentration in 50 mM Na^+^/K^+^ phosphate buffer, pH 5.8 supplemented with 100 mM NaCl, 1 mM TCEP, 100 µM DSS and 10 % D_2_O.

Circular dichroism spectra showed a shift of positive ellipticity from 274 nm for unstructured DNA (T_4_CAT) to 286 nm for dC-hairpin which was also accompanied by increase of molar ellipticity (**Extended Data Fig. S10A**). Presence of G-C base-pairs in the duplex part of dC-hairpin is evident from four singlets of four imino protons at 13-13.2 ppm in ^1^H NMR spectrum (**Extended Data Fig. S10B**). These data confirm that dC-hairpin is folded in solution and can be used as a scaffold to design A3A inhibitors by using nucleoside-based inhibitors of CDA instead of dC in the loop of dC-hairpin.

### Nuclear Magnetic Resonance (NMR) Spectroscopy and Mass Spectrometry of Hairpin DNA

^1^H, ^13^C, ^31^P NMR spectra were recorded on Bruker 500- and 700-MHz spectrometers, the latter with dual-channel cryoprobe. A representative spectrum in the imino region is shown in Figure S1B. NMR spectra were processed in TopSpin. High-resolution electrospray mass spectra were recorded on a Thermo Fisher Scientific Q Exactive Focus Hybrid Quadrupole-Orbitrap mass spectrometer. Ions generated by ESI were detected in positive ion mode for small molecules and negative ion mode for oligonucleotides. Total ion count (TIC) was recorded in centroid mode over the *m*/*z* range of 100-3,000 and analyzed using Thermo Fisher Xcalibur Qual Browser. Mass-spectrometric data on hairpin DNA are presented in **Table S1**.

### Isothermal Titration Calorimetry (ITC) of Interaction of A3A with Hairpin DNA

Desalted unmodified TTC-hairpin oligo was purchased (Integrated DNA Technologies) at 1 *μ*mol synthesis scale and dissolved in TE buffer (10 mM Tris/HCl pH 7.9, 1 mM EDTA) to give 10 mM solutions. ITC experiments were conducted at 25 °C using a MicroCal ITC200 (now Malvern Instruments) isothermal titration calorimeter. Protein A3A-E72A, which is a catalytically inactive variant, was dialyzed and diluted with ITC buffer to concentrations of about 50 *μ*M (ITC buffer: 50 mM Na^+^/K^+^ phosphate, pH 6.0, 50 mM NaCl, 50 mM choline acetate, 2.5 mM TCEP, 200 μM EDTA with 30 mg/mL bovine serum albumin; after preparation, this buffer was frozen and defrosted before the experiments) and titrated with dC oligonucleotides dialyzed against the above ITC buffer. The concentration ratio of oligonucleotide in the syringe to protein in the cell is generally 10:1 (for 1:1 binding). **Table S3** presents full analysis of ITC results; **Figures S12 and S13** show the titration curves and derived plots of enthalpy changes *versus* stoichiometry ratio A3A-E72A:hairpin DNA.

### Enzymology of A3A with Hairpin Substrates and Inhibitors

Hairpins as substrates and inhibitors were analyzed as previously described.^33^ In short, wild type A3A was used to compare linear DNA (A_2_T_2_**C**A_4_) and dC-hairpin (T(GC)_2_TT**C**(GC)_2_T, bold C is deaminated) at 500 *μ*M in the NMR-based assay (pH 7.4, 50 mM Na^+^/K^+^ phosphate buffer, supplemented with 100 mM NaCl, 1 mM TCEP, 100 µM sodium trimethylsilylpropanesulfonate (DSS) and 10 % D_2_O; enzyme concentration: 140 nM, 20 °C and 150 mM salt concentration. The NMR-based assay yields the initial velocity of deamination of various ssDNA substrates, including the modified ones,^30^ in the presence of A3 enzymes. Consequently, the Michaelis–Menten kinetic model is used to characterize substrates and inhibitors of A3. Moreover, use of dC-containing hairpin as a substrate of A3A allowed us to use a global regression analysis of the kinetic data over the entire time course of the reaction using Lambert’s W function (integrated form of the Michaelis-Menten equation).

The course of the reaction was followed by ^1^H NMR until the substrate was consumed (28 hours). Subsequently the amount of substrate or product at each time point was calculated by integrating the decreasing substrate peak at 7.752 ppm (singlet) or the increasing product peak at 5.726 ppm (doublet) and calibrated by the area of DSS standard peak at 0.0 ppm. Using the known concentration of the standard, the peak was converted to a corresponding substrate concentration. The time at which each spectrum was recorded as a difference to the first spectrum was used as the time passed. The product or substrate concentration *versus* the time of reaction was plotted and fitted using the integrated form of the Michaelis-Menten equation:

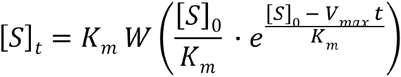

where *W* is Lambert’s W function, [*S*]_*t*_ is the substrate concentration at specific time, [*S*]_0_ is the initial substrate concentration, *V*_max_ and *K*_m_ are the Michaelis-Menten constants and *t* is the time. The two Michaelis-Menten constants, *k*_cat_ and *K*_m_, the initial substrate concentration and an offset which corrects for the integration baseline in the NMR spectra were fitted using Lambert’s *W* function in Gnuplot.

By varying the concentration of an inhibitor, the plots of observed *K*_m_ versus inhibitor concentration were obtained, fitted with a linear function (*f*(*x*) = *a* + *b x*) and *K*_i_ values were calculated as *a*/*b*, with error propagation as described previously (**Figure S11**).^30^

### Evaluation of Stability of PS-hairpins against Enzymatic Digestion

Separately TTC-hairpin, PS-TTFdZ-hairpin, and 3*′*-fluorescein-labelled PS-TTdZ-hairpin-FAM and PS-TTdZ-hairpin-FAM (each 15 *μ*M in 50 mM Tris-HCl buffer, 10 mM MgCl_2_, pH 8.0, 37 °C) were treated with snake venom phosphodiesterase (phosphodiesterase I, Sigma, 32 mU/mL). The percent degradation over time (0-360 minutes) was monitored by anion-exchange chromatography for the indicated times at 37 °C.

### DNA Deaminase Activity Assay

HEK 293T (RRID: CVCL_0063) cells (ATCC, USA) were maintained in RPMI-1640 (#SH30027.01, Cytiva, USA) and supplemented with 10% fetal bovine serum (#10437028, Gibco, ThermoFisher, USA) at at 37 °C with 5% CO_2_ in a humidified atmosphere. The deaminase activity was performed as previously described^55^. Lysate was placed on ice and immediately used for the deaminase assay. Inhibitor oligos (and controls) were heated to 80°C for 5 minutes, and then cooled to RT to induce hairpin formation. They were then incubated at varying concentrations in 5 µL with 10 µL of cell lysate at 37°C for 15 minutes to promote binding of oligos to A3A. To this, 5 µL of: 0.25 µL RNAse A, 800 nM fluorescent oligo, 10x UDG buffer, and 0.25 µL UDG were added to each sample for a total volume of 20 µL and incubated at 37°C for 1 hour. The fluorescently-labeled oligo has the following sequence: (5′-ATTATTATTATT**<u>C</u>**GAATGGATTTATTTATTTATTTATTTATTT-fluorescein-3′). The samples were run on a 15% Urea-TBE acrylamide gel, imaged with a Typhoon FLA-7000 imager (GE Healthcare), and then quantified using ImageQuant (Cytiva, USA).

### Cellular Uptake and Localization of DNA Oligomers

MCF7 (RRID: CVCL_0031) cells (ATCC, USA) were maintained in DMEM (Gibco, ThermoFisher Scientific, USA) supplemented with 1% penicillin/streptomycin (Gibco), and 10% fetal bovine serum (Gibco) at 37 °C with 5% CO_2_ in a humidified atmosphere.

Briefly, MCF7 cells were transfected with FAM-labelled hairpins using X-tremeGENE 9 DNA Transfection Reagent (Roche, USA; 1.0 µL). After 16 hours, the cells were washed twice with phosphate-buffered saline (PBS) containing MgCl_2_ and CaCl_2_, fixed in 4% paraformaldehyde/PBS for 15 min at room temperature (RT), and then washed with PBS. Cells were then stained with Hoechst 33342 before imaging on an Zeiss LSM 900 Scanning Confocal Microscope using an oil-immersion 63 objective lens (NA 1.4). Laser excitation wavelengths and collection ranges appropriate to the fluorophores of each sample were used to detect the emission spectra of the specific combination of Hoechst 33342 (excitation at 405 nm, emission monitored at 410–530 nm) and FAM 555 (excitation at 555 nm, emission monitored at 565–600 nm). All images were digitally processed for presentation with ImageJ (Rasband, W. 2014. ImageJ. U.S. National Institutes of Health, Bethesda, MD). Confocal microscopy images of cellular uptake of inhibitors are provided in **Figure 3A-C)** and Figures **S14-S18**.

### Restriction of A3A Activity *in Cellulo*

HEK 293T cells stably transduced with the live-cell deaminase reporter have been previously reported^45-46^. Semi-confluent cells in a 24-well TPP plate (#Z707791, Millipore Sigma, Merck, DE) were transfected with the following: pcDNA3.1-A3A-HA (20 ng), LTR-gRNA-Cas9n-UGI-NLS-Puro-LTR (400 ng), and varying concentrations of inhibitor (or control) oligos using TransIT-LT1 (Mirus Bio) as the transfection reagent. The plate was imaged using an Incucyte (Sartorius, USA) over the course of 68 hours. Live-cell images of orange, green, and phase image channels were captured with a 4x objective every four hours after the initial transfection with five images per well in a fixed grid. mCherry- and GFP-positive cells were identified with internal cellular analysis software. The rolling slope for each concentration between 30 and 50 hours was calculated using GraphPad Prism (Dotmatics) software (first derivative between two time points) and represented as an average in **Fig. 3d**. After 68 hours, cells were collected and prepared for immunoblots.

### Immunoblots

Immunoblots were prepared as previously described^46^. Samples were separated by either a 4-20% Criterion TGX Precast gel (#5671095, Bio-Rad, USA) or 4-15% Mini PROTEAN TGC gel (#4561084, Bio-Rad, USA), and then transferred to nitrocellulose membranes (#1620112, Bio-Rad, USA). Primary antibodies include mouse anti-Tubulin (#T5168, 1:10000, Sigma-Aldrich), rabbit anti-HA (#3724S, 1:2500, Cell Signaling), and rabbit anti-Cas9 (#ab189380, 1:5000, Abcam). Secondary antibodies used were goat anti-rabbit IRdye800 (LI-COR, #925-32211, 1:10000) and goat anti-mouse IRdye680 (LI-COR, #926-69020, 1:10000). Raw immunoblots are included in Supplementary Information, **Fig. S1**.

## DATA AVAILABILITY

Coordinates and structure factors are available with the following PDB codes: 8FIM (A3A-E72A with TTC-hairpin DNA substrate), 8FIL (Zn^2+^-free A3A-E72A with TTC-hairpin DNA substrate), 8FIK (A3A-E72A with ATTC-hairpin DNA substrate), 8FIJ (wildtype A3A with TTFdZ-hairpin inhibitor form 2), 8FII (wildtype A3A with TTFdZ-hairpin inhibitor form 1).

## ACKNOWLEDGEMENTS

Financial support of the Health Research Council of New Zealand in partnership with Breast Cancer Research (grant 20/1355) and Kiwi Innovation Network with Massey Ventures Limited (grant MU002391), as well as the support of the School of Natural Sciences, Massey University is gratefully acknowledged. Access to the MX2 Beam Line of the Australian Synchrotron was facilitated by the NZ Synchrotron Group Ltd financially supported by the Ministry of Business Innovation and Enterprise (MBIE) and a consortium of universities, including Massey University. We are grateful to beam-line scientist Dr Alan Riboldi-Tunnicliffe for assistance with remote access to the Australian Synchrotron during Covid19 lockdown. We thank Dr Patrick J. B. Edwards and Mr David Lun for their assistance with use of the NMR and mass spectrometry facilities at Massey University. We thank B. Moriarity for contributing to the mentorship of AER. Cancer studies in the Harris lab are supported by NCI P01-CA234228 and a CPRIT Established Investigator Recruitment Award. RSH is an Investigator of the Howard Hughes Medical Institute, a CPRIT Scholar, and the Ewing Halsell President’s Council Distinguished Chair at University of Texas Health San Antonio.

## AUTHOR CONTRIBUTIONS

VVF, EH, TKH and RSH designed experiments, supervised the research and analyzed data; SH, HMK, YS, GBJ, MB, JF, TKH, and AER performed experiments; GBJ wrote the first draft and VVF, EH, TMK, AER, and RSH added sections and edited the manuscript.

## ADDITIONAL INFORMATION

Supplementary information is available and includes additional information on the structures, table of isothermal titration calorimetry data, table of kinetic results, and original, uncropped immunoblots.

**Extended Data Fig. 1.**
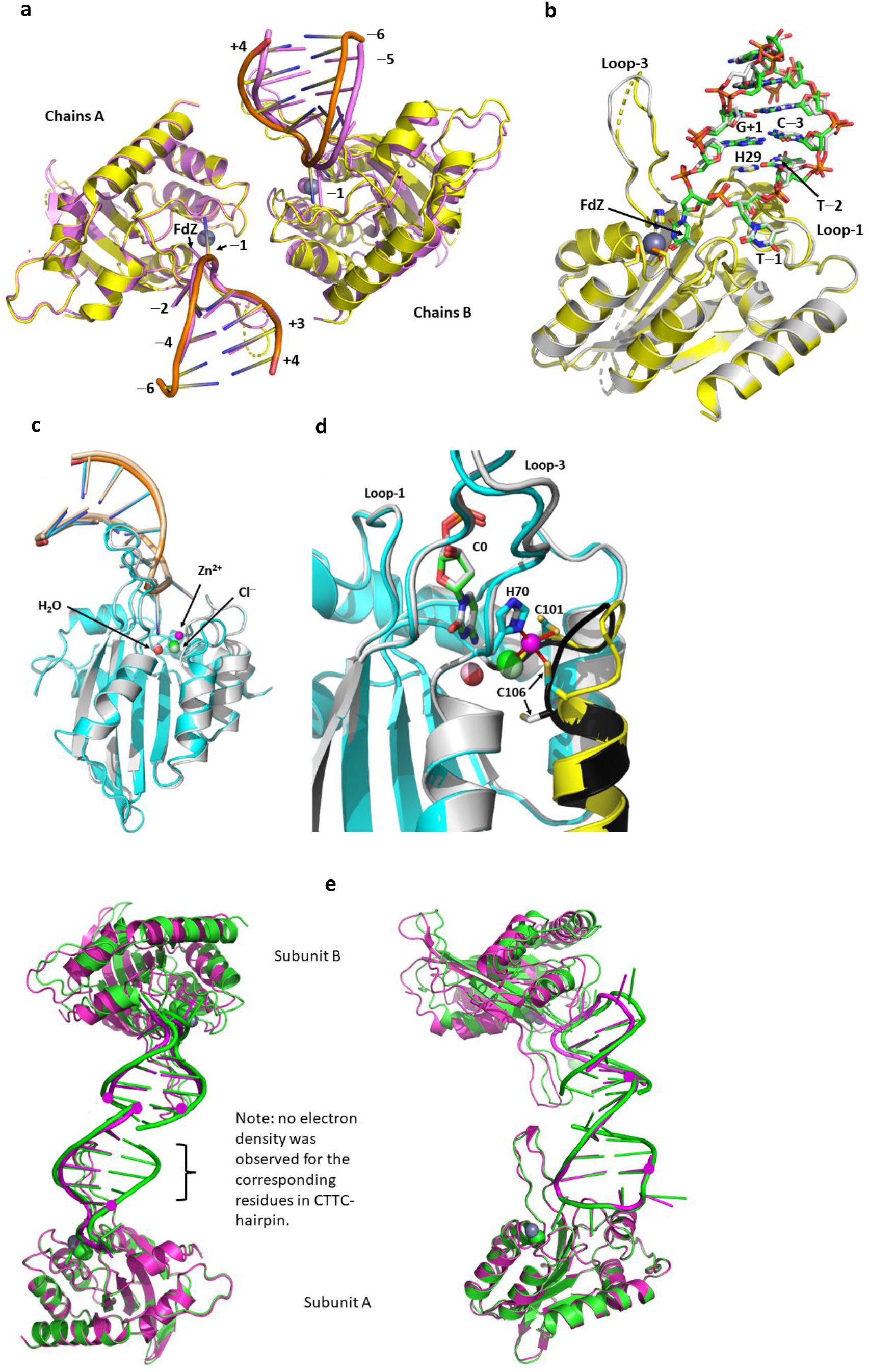
Superpositions of structures of wildtype A3A and its inactive EE72A mutant in complexes with hairpin DNA. **a**, Superposition of subunits A of wildtype A3A complexed with TTFdZ-hairpin. The 2.94 Å resolution structure is shown in yellow (protein) and orange (hairpin); the 2.80 Å structure is shown in light purple. Note displacement and small rotation of chain B for the latter relative to the former. In general loops are better defined in chain B for both structures, whereas for the former (2.94 Å resolution) structure, the hairpin is much better defined. The Zn^2+^ is shown as grey and light purple spheres, respectively. **b**, Superposition of subunit B (carbon atoms in grey-blue) onto subunit A (carbon atoms of protein in yellow, of hairpin in green)) for the 2.94 Å resolution structure (RMSD = 0.47Å). Key active-site residues (His70, Cys101, Cys106, Glu72 and His29) are shown in stick form. Zn^2+^ is shown as a grey sphere. Note: tight NCS restraints are placed on torsion angles of the pair of subunits. The small differences in conformations of the hairpin are not considered significant. **c**, Superposition of structures of A3A-E72A complexed with TTC-hairpin where the Zn^2+^-containing structure is in cyan and the Zn^2+^-free structure is in grey. Diagram showing similarity in overall structure of protein and hairpin DNA. For the Zn^2+^-free structure, the Cl^−^ is coloured light-green, the water light-red and the hairpin wheat. **d**, Diagram of the active site showing reconfiguration of the loop bearing Cys101 and Cys106 in the absence of Zn^2+^. The loop is highlighted in yellow for Zn^+2^-containing structure and black for the Zn^2+^-free structure. Cytosine C^0^ is shown in sticks (green for Zn^2+^-containing structure; grey for Zn^2+^-free structure). Note small displacements of active site water, chloride ion and ligands His70 and Cys101, but large displacement of Cys106 on loss of Zn^2+^. **e**, Superposition of A subunits of complexes of A3A-E72A with CTTC-hairpin (magenta) onto the complex with ATTC-hairpin (green). The apparent stem bases for CTTC-hairpin are marked by magenta spheres. Note that for CTTC-hairpin, one hairpin adopts the expected threading shown in **Extended Data Table 1**, but the other hairpin appears to be threaded differently, placing C^−3^ (of **Extended Data Table 1** threading) into the active site of A3A. Left frame: the view is approximately down the two-fold NCS element. Right frame: relative to left frame, the pair of molecules has been rotated about horizontal (∼30°) and vertical (∼90°) axes.

**Extended Data Fig. 2.**
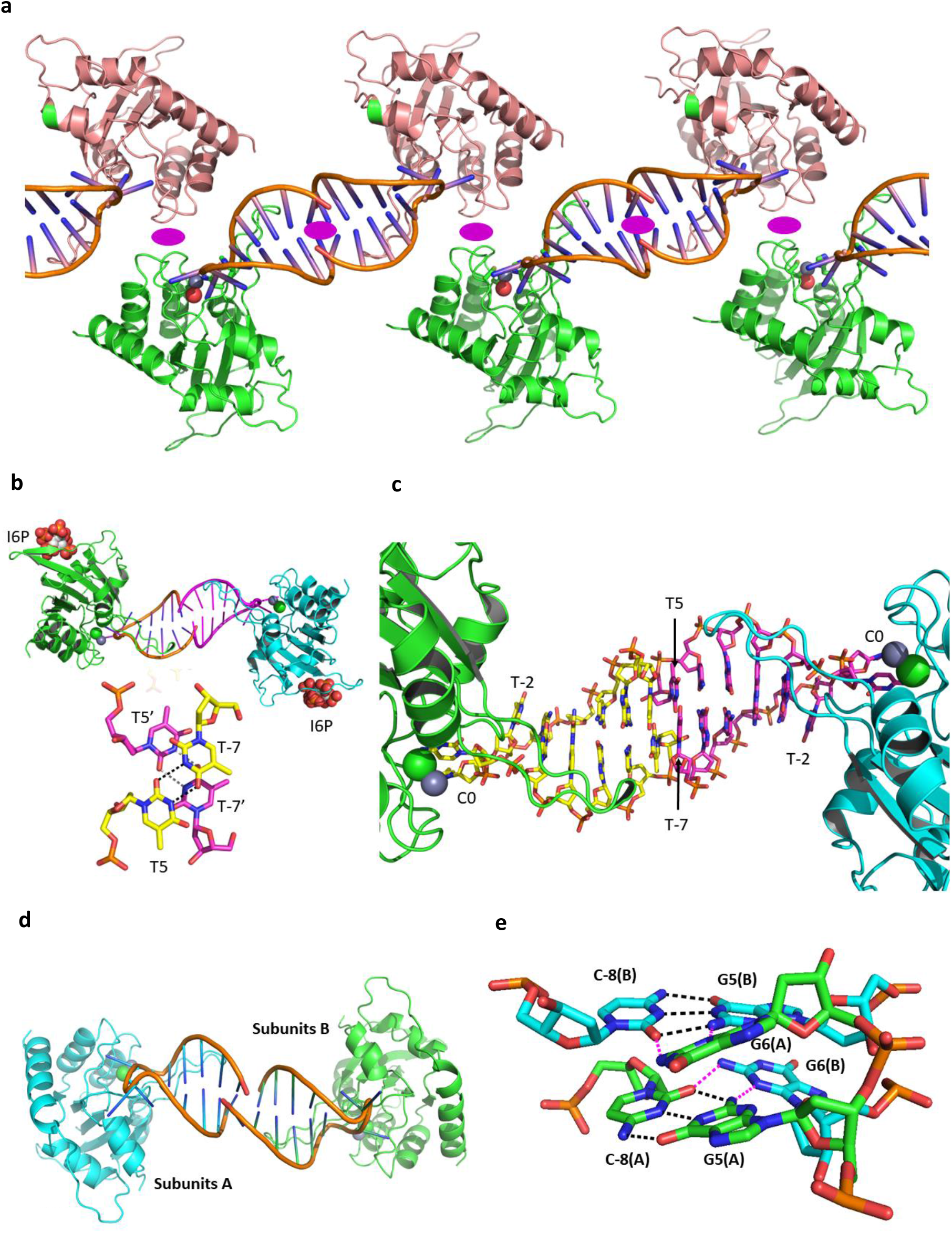
Schematic diagram showing the general shared packing of A3A in the unit cell and the packing of the stem bases for different hairpins. **a**, In this A3A-E72A with TTC-hairpin and other structures presented here, there are two molecules in the asymmetric unit, arranged such that the base of one hairpin stacks with the base of the hairpin of the other molecule with approximate non-crystallographic two-fold symmetry. The NCS two-fold elements are shown by purple ellipses. **b**, Cartoon representation of a pair of molecules for A3A-E72A in complex with TTC-hairpin. Inositol hexaphosphate (phytic acid, I6P; shown as spheres) mediates intermolecular interactions of A3A-E72A. Stick representation detailing the interaction between the bases of the stems. **c**, Stick representation showing the base pairing and projection of the TTC loop into A3A. **d**, Cartoon representation of A3A-E72A in complex with ATTC hairpin showing the interaction of the base of the stem of ATTC-hairpin of one molecule with the base of the stem of another molecule. Cartoon representation of A3A-E72A in complex with ATTC-hairpin. The two-fold NCS is approximately out of the plane. **e**, Stick representation showing detail of the hydrogen bonding of the penultimate C^−8^G^5^ pair (black dashes) and the hydrogen bonding of the final G^6^ of one hairpin with the NCS-related C^−8^G^5^ hairpin (magenta dashes).

**Extended Data Fig. 3.**
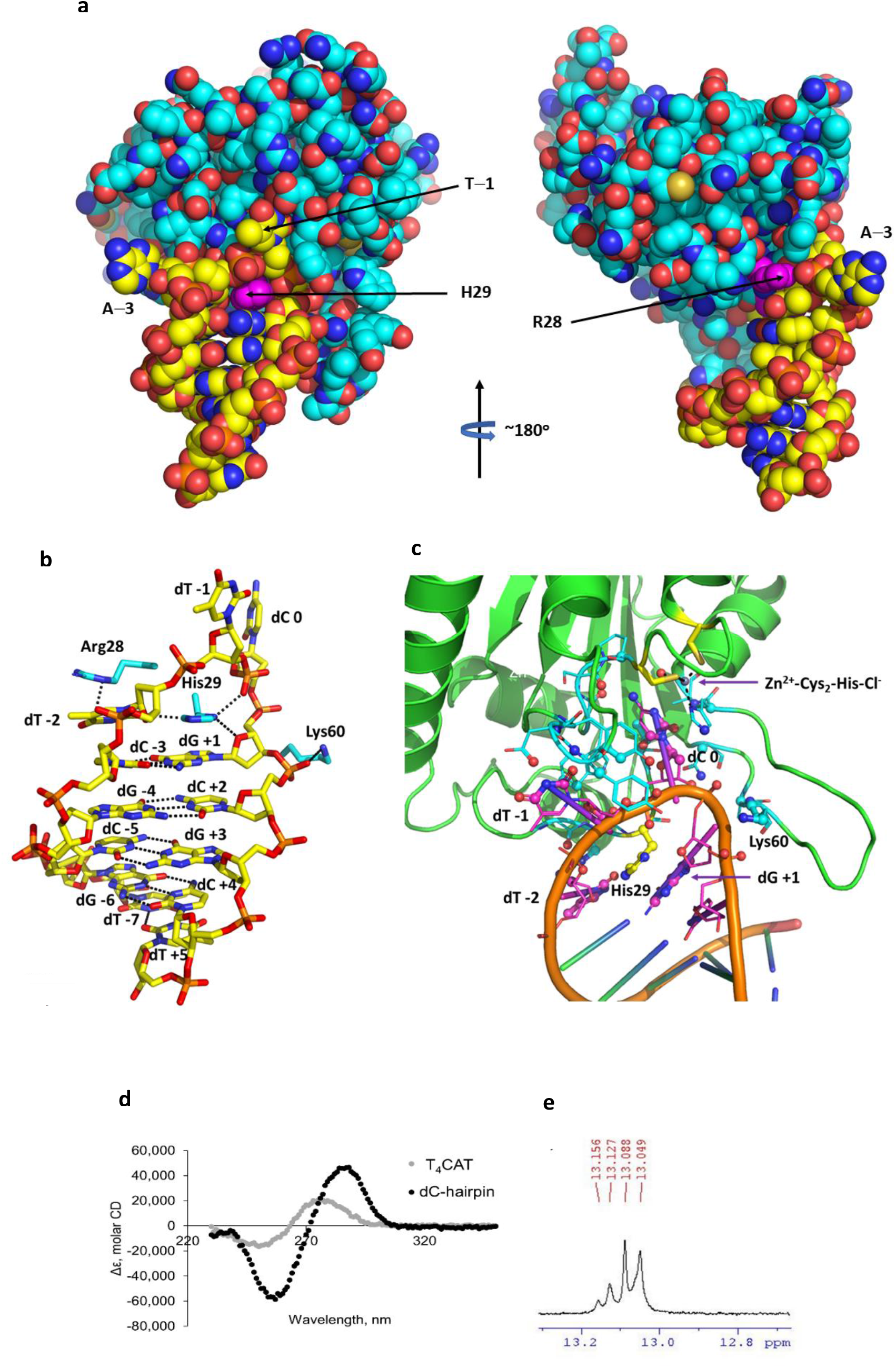
Interactions of ATTC-hairpin with A3A-E72A and evidence for hairpin structure in solution. **a**, Space-filling diagram of substrate/inhibitor binding to A3A, showing crucial role of His29 in the binding of substrate/inhibitor to A3A. His29 and Arg28 are highlighted in magenta; protein carbon atoms are shown in cyan and those for the ATTC-hairpin in complex with A3A-E72A; all other atoms in standard CPK colors. **b**, Base-pairing in the stem of ATTC-hairpin and key interactions of the hairpin with Arg28, His29 (especially) and Lys60 (additional to interactions of dC and dT^−1^ with A3A). Similar interactions occur for ATTC- and CTTC-hairpins (less precisely) with A3A-E72A and for TTFdZ-hairpin with wildtype A3A (also less precisely). **c**, Protein residues in contact with the oligo are shown in cyan for carbon atoms and oligo residues in contact with the protein are shown in magenta for carbon atoms. Atoms in contact (<3.5 Å) are shown as spheres (∼40 protein and ∼40 oligo atoms; ∼10 residues and 4 nucleotides). **d**, Circular dichroism spectra of linear T_4_CAT and TTC-hairpin (no A3A). **e**, ^1^H NMR spectrum of TTC-hairpin in the imino region. Both spectra are consistent with formation of a hairpin structure in solution.

**Extended Data Fig. 4.**
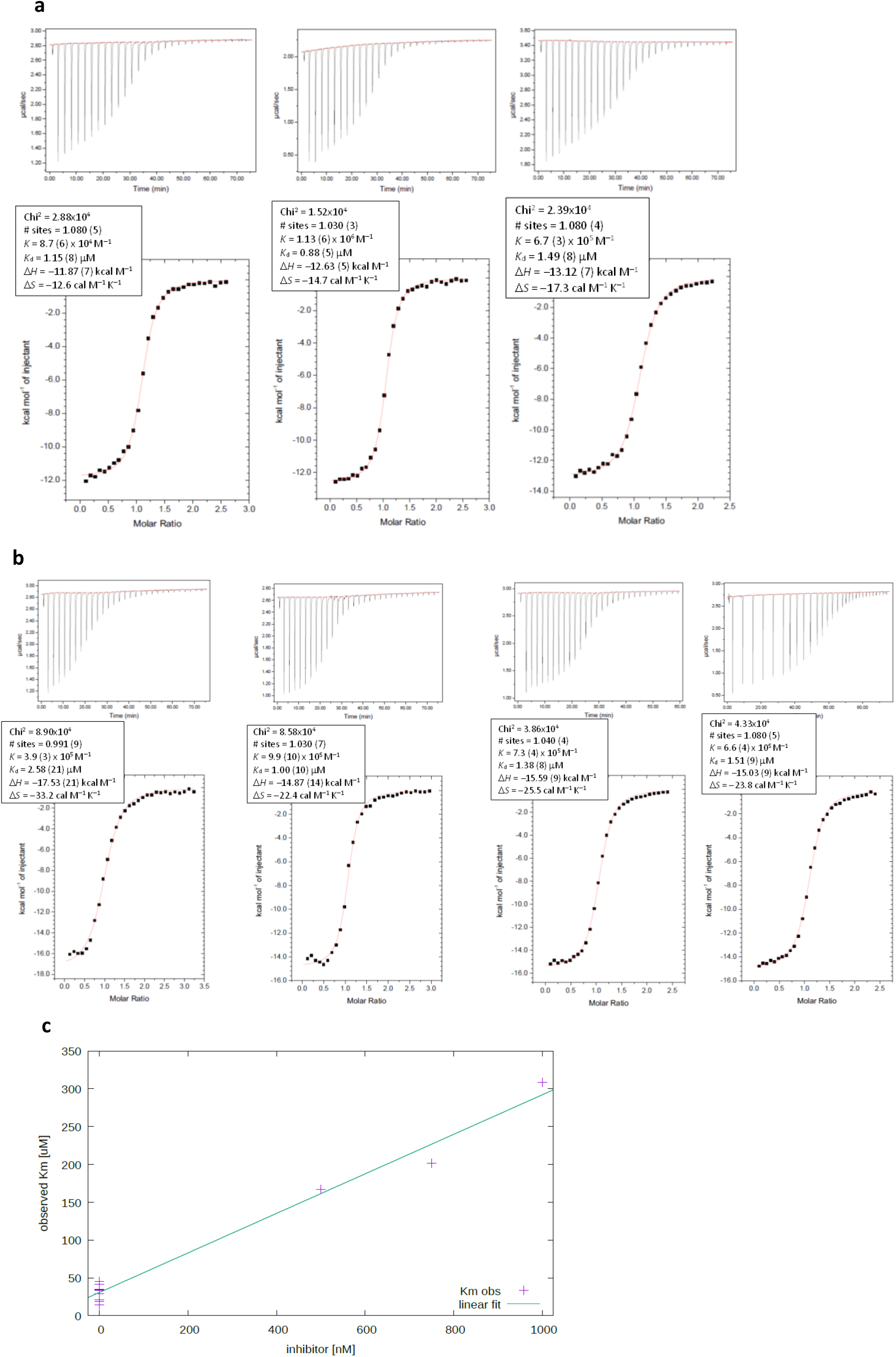
Isothermal titration calorimetry (ITC) data and inhibition data. **a**, ITC titration of A3A-E72A with TTC-hairpin (3 replicates). See Methods for experimental conditions. **b**, ITC titration of A3A-E72A with PS-TTC-hairpin (4 replicates). See Methods for experimental conditions. **c**, Plot of apparent *K*_m_ *versus* inhibitor concentration for reaction of TTC-hairpin with wildtype A3A in the presence TTFdZ-hairpin inhibitor. The value of *K*_i_ is obtained from the ratio of slope/intercept of this graph.

**Extended Data Fig. 5.**
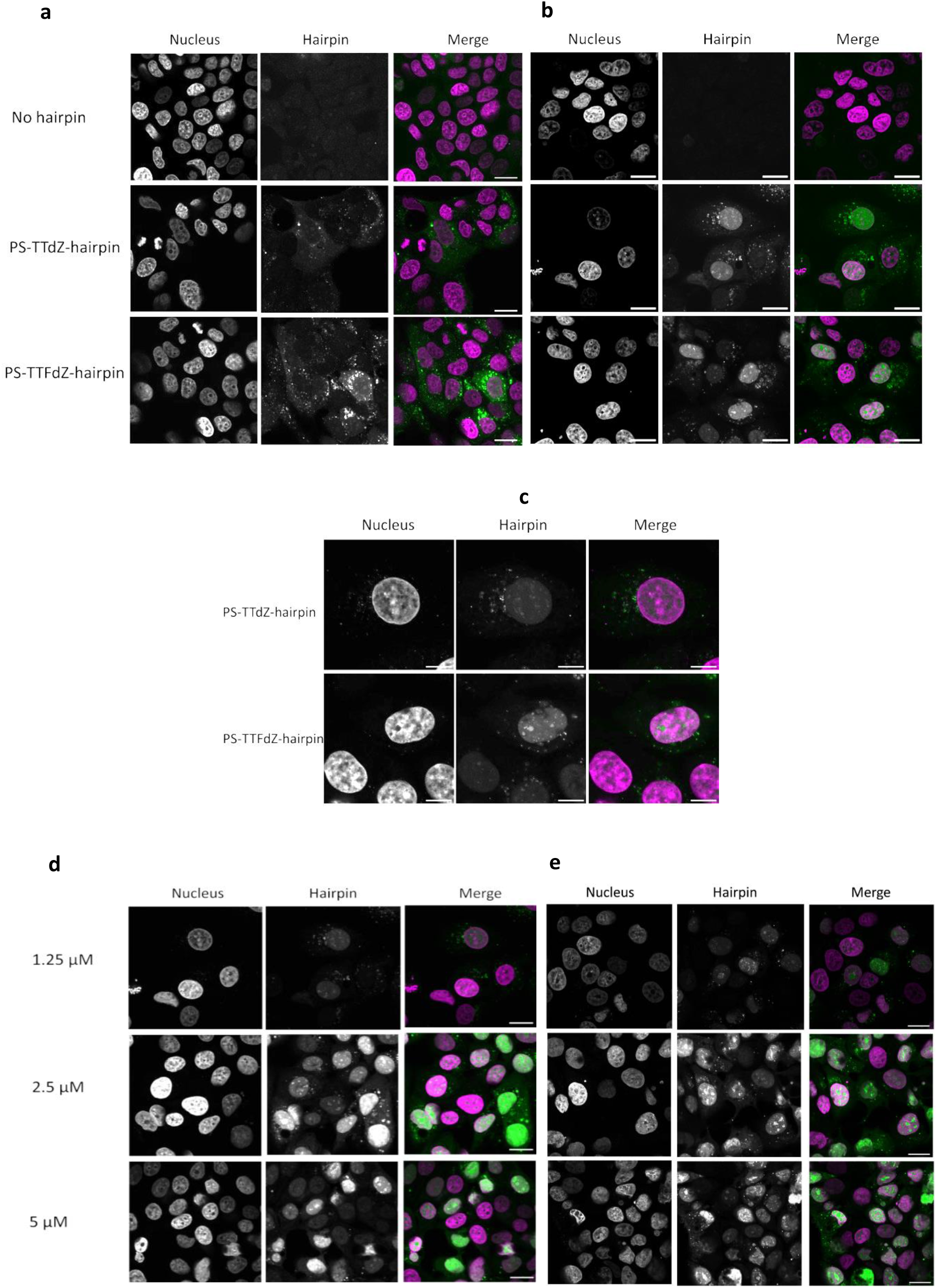
Representative confocal microscopy images of MCF-7 cells incubated with FAM-labelled phosphorothioated- (PS-) modified hairpins at varying concentrations (0-20 μM). Hoechst 3342 is used for staining of the nucleus and the hairpin is labelled with fluorescein (FAM) at the 3*′* end. Individual panels, nucleus/Hoechst 33342 (left) and oligo/FAM (middle) are shown for each section, along with merge where pseudo-coloured panels (right) are overlaid, nucleus (magenta) and oligo-FAM (green). Except where indicated otherwise, the scale bars are 20 μm. **a**, Representative images of MCF-7 cells showing absence of uptake of 20 *μ*M PS-modified FAM-labelled hairpins (no oligo (top panels) PS-TTdZ-(middle panels) and PS-TTFdZ-hairpin (bottom panels)) *in the absence of transfection reagent after 16 hours*. **b**, Representative images of MCF-7 cells incubated with no hairpin (upper panels) or 1.25 μM of PS-TTdZ-(middle panels) or PS-TTFdZ-FdZ (lower panels) hairpins *in the presence of X-tremeGENE 9 DNA Transfection Reagent after 16 hours*. **c**, Zoomed images of cells shown in (**b**). Scale bars, 10 *μ*m. **d**, Representative images of MCF-7 cells incubated with PS-TTdZ-hairpin at 1.25 μM (upper panels), 2.5 μM (middle panels), 5 μM (lower panels) in the presence of transfection reagent (Xtreme GENE™ HP) after incubation for 16 hours. **e**, Representative images of MCF-7 cells incubated with PS-TTFdZ-hairpin at 1.25 μM (upper panels), 2.5 μM (middle panels), 5 μM (lower panels) in the presence of transfection reagent (Xtreme GENE™ HP) after incubation for 16 hours.

**Extended Data Fig. 6.**
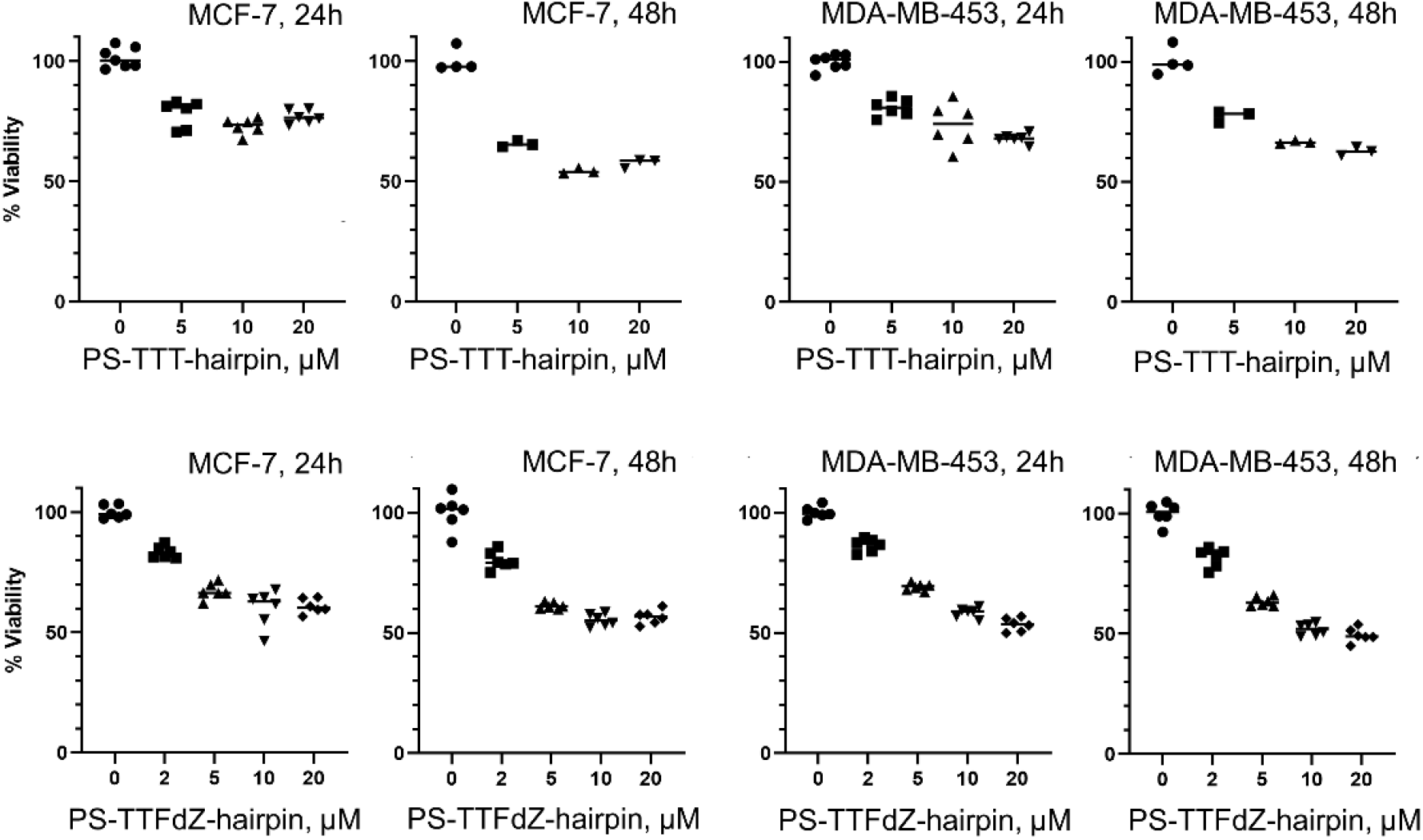
Viability (%) of MCF-7 and MDA-MB-453 cells at various concentrations PS-modified hairpins. PS-TTT-hairpin (upper panels) and PS-TTFdZ-hairpin (lower panels) at 24 h and 48 h, as indicated on the Figure. Data are normalized to average number of cells in the presence of transfection reagent only. Data were collected from three individual experiments with biological replicates except for the PS-TTT-hairpin at 48 h.

**Extended Data Fig. 7.**
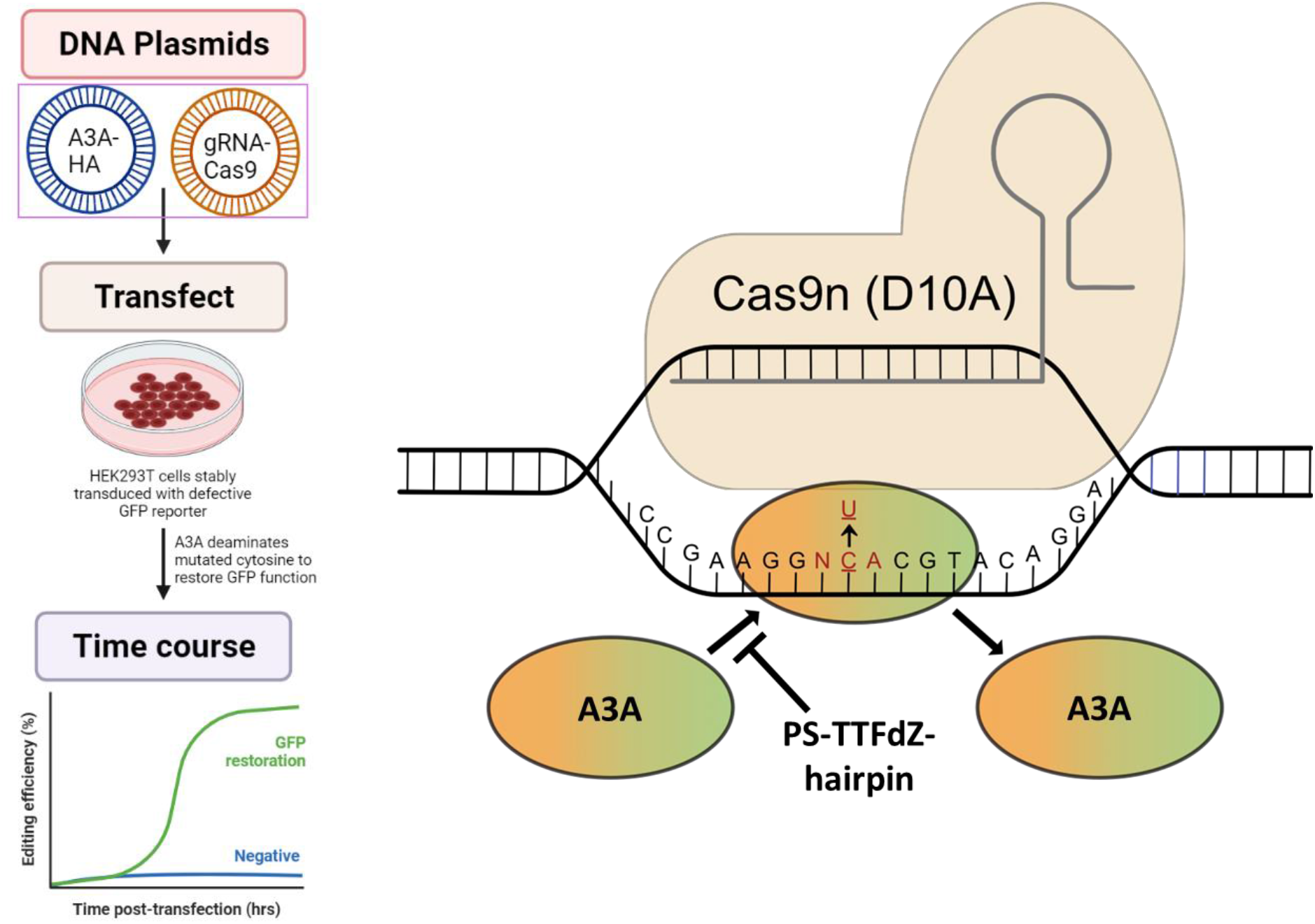
The transfection of HEK293 and expression of A3A. Left frame summarizes the transfection and monitoring of GFPO fluorescence procedure. The right frame shows that A3A, in absence of inhibitor, corrects *Gfp* to express functional GFP. The Cas9n is directed to the *Gfp* 5’-T<u>C</u>A site-of-interest using a complementary gRNA and opens an R-loop that A3A (here unlinked to Cas9n) edits to 5’-T<u>T</u>A to restore GFP functionality. PS-TTFdZ-hairpin inhibits this DNA-editing by A3A. Left panel schematic was designed using BioRender.

**Extended Data Fig. 8.**
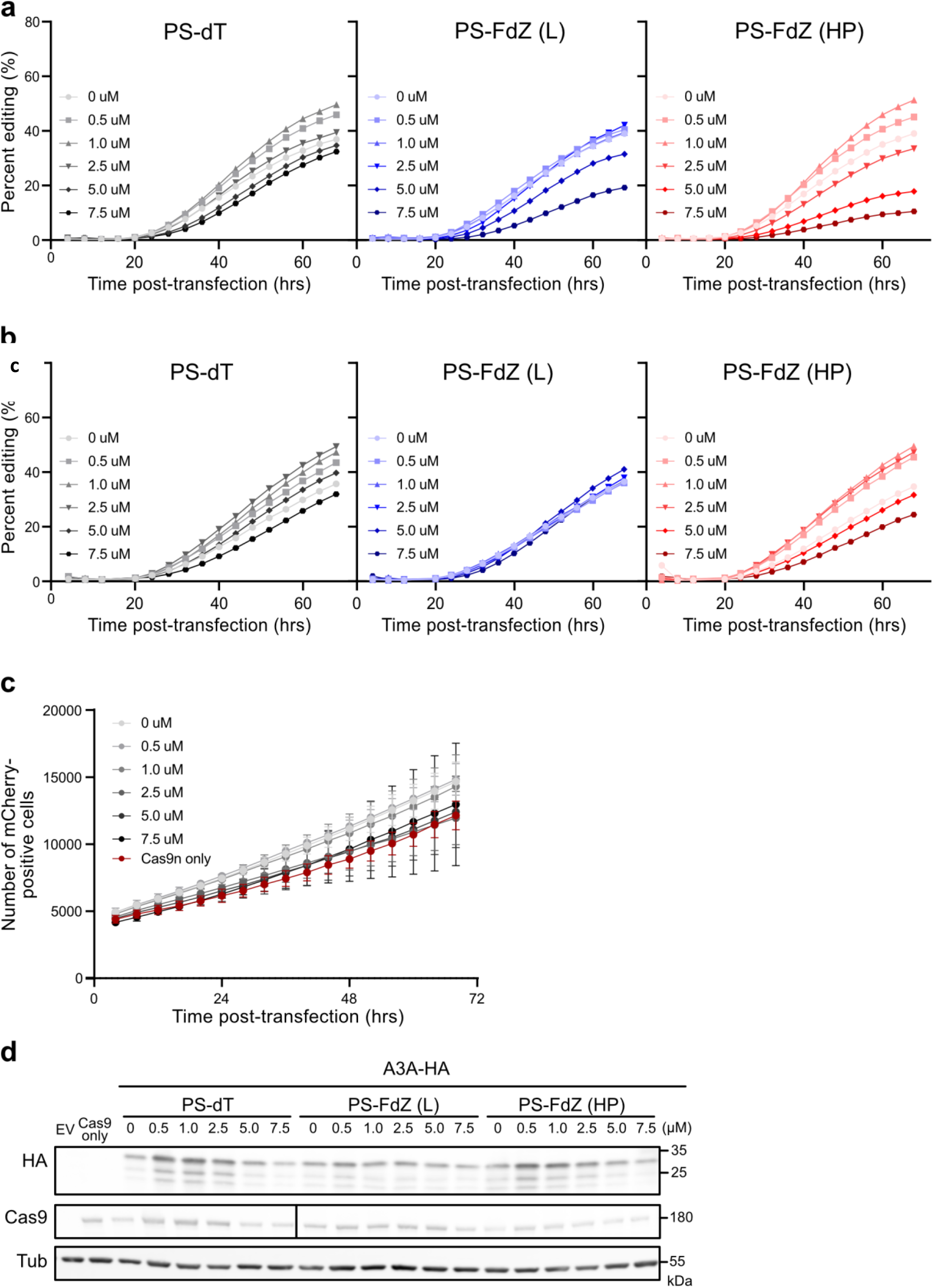
Replicates of *in cellulo* assay of A3A-editing activity by linear and hairpin DNA. PS-TTFdZ-linear is denoted FdZ (L); PS-TTT-hairpin (as control) is denoted dT; and PS-TTFdZ-hairpin is denoted FdZ HP for a total of three biologically independent replicates (see Fig. 3). **b**, Representative example of mCherry cell count over time during live cell assay. Error bars depict mean of cell count between the three oligos tested [dT, FdZ (L), and FdZ (HP)] at each concentration. The red line represents the Cas9-only control (two technical replicates). **c**, Representative immunoblots of samples shown in **Fig. 3d**. Antibodies used are detailed Methods. The black bar in the Cas9 panel was used to help correctly line up bands that were skewed due to curving of blot. Raw blots are shown in Supplementary Information, **Fig. S1**.

**Extended Data Table 1.** List of oligos synthesized

| Name | DNA sequence,a) 5'→3' | ε260 (L·mol<sup>-1</sup>·cm<sup>-1</sup>) | Retention time (min)b) | ESI-MS [Da] found/calculated |
| --- | --- | --- | --- | --- |
| PS-TTC-hairpin a) | GpsCpsGpsCpsTpsTpsCpsGpsCpsGpsC | - | - | - |
| TTFdZ-hairpin | GCGCTT**FdZ**GCGC | 94800 | 14.3 | 3311.5/3311.6 |
| TTFdZ-hairpin<br>(used for crystallization) | TGCGCTT**FdZ**GCGCT | 112200 | 14.1 | 3919.6/3918.7 |
| PS-TTFdZ-hairpin | GpsCpsGpsCpsTpsTps**FdZ**psGpsCpsGpsC | 94800 | 27.6<sup>c)</sup> | 3472.3/3472.7 |
| PS-TTdZ-hairpin | GpsCpsGpsCpsTpsTps**dZ**psGpsCpsGpsC | 94800 | 17.7 | 3454.3/3454.7 |
| PS-TTT-hairpin | GpsCpsGpsCpsTpsTpsTpsGpsCpsGpsC | 101700 | 17.8 | 3482.4/3482.3 |
| PS-FdZ-linear | ApsTpsTps**FdZ**psApsTpsTpsT | 84800 | 19.3 | 2810.2/2802.3 |
| PS-dZ-linear | ApsTpsTps**dZ**psApsTpsTpsT | 84800 | 19.2 | 2791.2/2790.3 |
| PS-TTFdZ-hairpin- FAM | GpsCpsGpsCpsTpsTps**FdZ**psGpsCpsGpsCps/FAM | 115760 | 29.6<sup>c)</sup> | 4138.4/4139.3 |
| PS-TTdZ-hairpin-FAM | GpsCpsGpsCpsTpsTps**dZ**psGpsCpsGpsCps/FAM | 115960 | 30.2<sup>c)</sup> | 4120.4/4119.3 |

a) PS refers to phosphorothioate linkage. This particular oligo was used as validated and supplied by IDT.
b) Reverse-phase HPLC, except where otherwise flagged.
c) Ion-exchange HPLC.

**Extended Data Table 2.**
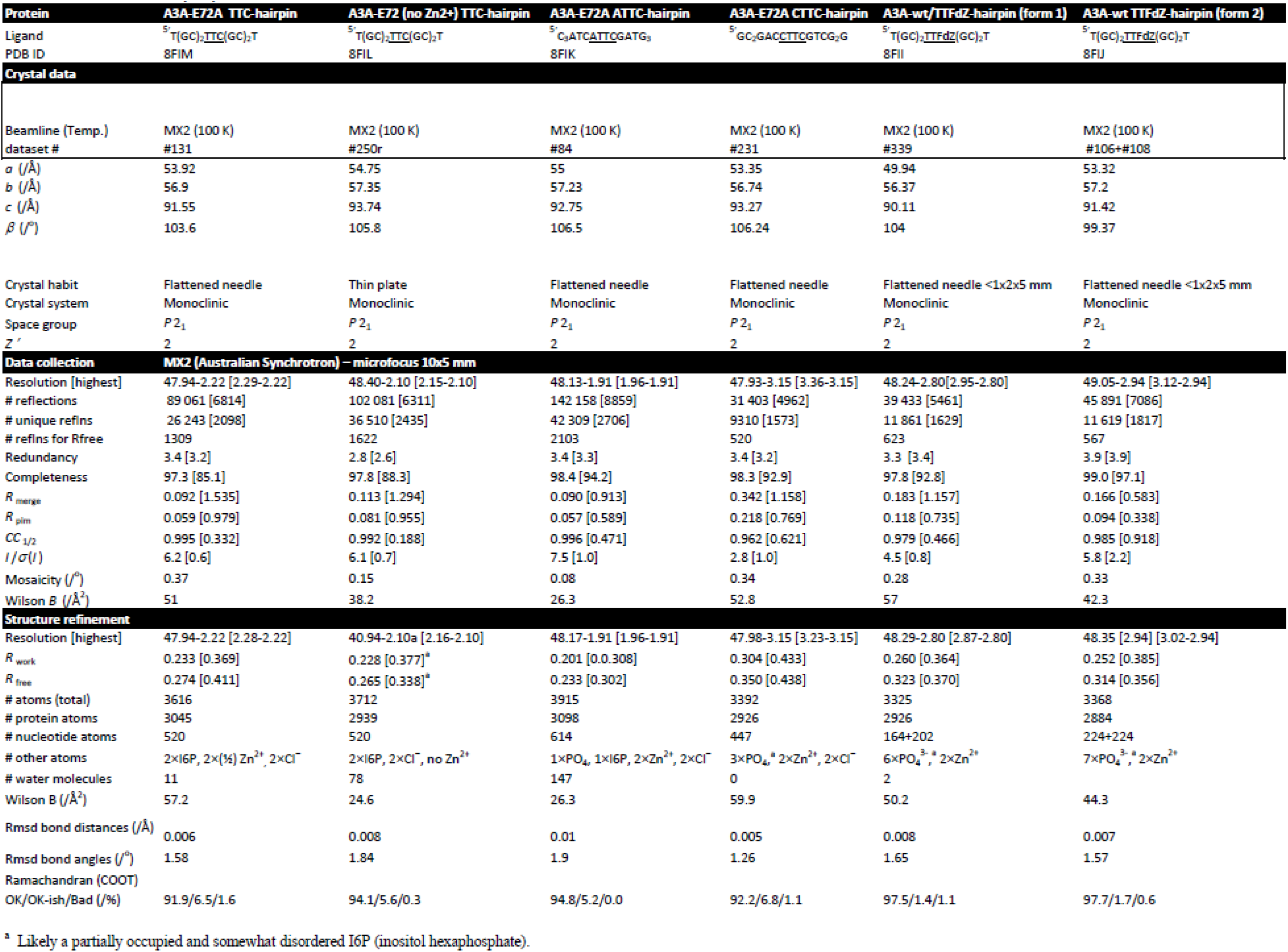
Crystal data, data collection and refinement details

## REFERENCES

1. Harris, R. S.; Bishop, K. N.; Sheehy, A. M.; Craig, H. M.; Petersen-Mahrt, S. K.; Watt, I. N.; Neuberger, M. S.; Malim, M. H., DNA deamination mediates innate immunity to retroviral infection. Cell 2003, 113 (6), 803–809.

2. Harris, R. S.; Liddament, M. T., Retroviral restriction by APOBEC proteins. Nature Reviews Immunology 2004, 4 (11), 868–877.

3. Izumi, T.; Shirakawa, K.; Takaori-Kondo, A., Cytidine deaminases as a weapon against retroviruses and a new target for antiviral therapy. Mini-Rev. Med. Chem. 2008, 8 (3), 231–238.

4. Mangeat, B.; Turelli, P.; Caron, G.; Friedli, M.; Perrin, L.; Trono, D., Broad antiretroviral defence by human APOBEC3G through lethal editing of nascent reverse transcripts. Nature 2003, 424 (6944), 99–103.

5. Green, A. M.; Weitzman, M. D., The spectrum of APOBEC3 activity: From anti-viral agents to anti-cancer opportunities. DNA Repair (Amst) 2019, 83, 102700.

6. Swanton, C.; McGranahan, N.; Starrett, G. J.; Harris, R. S., APOBEC enzymes: mutagenic fuel for cancer evolution and heterogeneity. Cancer discovery 2015, 5 (7), 704–712.

7. Alexandrov, L. B.; Nik-Zainal, S.; Wedge, D. C.; Aparicio, S. A.; Behjati, S.; Biankin, A. V.; Bignell, G. R.; Bolli, N.; Borg, A.; Børresen-Dale, A. L.; Boyault, S.; Burkhardt, B.; Butler, A. P.; Caldas, C.; Davies, H. R.; Desmedt, C.; Eils, R.; Eyfjörd, J. E.; Foekens, J. A.; Greaves, M.; Hosoda, F.; Hutter, B.; Ilicic, T.; Imbeaud, S.; Imielinski, M.; Jäger, N.; Jones, D. T.; Jones, D.; Knappskog, S.; Kool, M.; Lakhani, S. R.; López-Otín, C.; Martin, S.; Munshi, N. C.; Nakamura, H.; Northcott, P. A.; Pajic, M.; Papaemmanuil, E.; Paradiso, A.; Pearson, J. V.; Puente, X. S.; Raine, K.; Ramakrishna, M.; Richardson, A. L.; Richter, J.; Rosenstiel, P.; Schlesner, M.; Schumacher, T. N.; Span, P. N.; Teague, J. W.; Totoki, Y.; Tutt, A. N.; Valdés-Mas, R.; van Buuren, M. M.; van ‘t Veer, L.; Vincent-Salomon, A.; Waddell, N.; Yates, L. R.; Zucman-Rossi, J.; Futreal, P. A.; McDermott, U.; Lichter, P.; Meyerson, M.; Grimmond, S. M.; Siebert, R.; Campo, E.; Shibata, T.; Pfister, S. M.; Campbell, P. J.; Stratton, M. R., Signatures of mutational processes in human cancer. Nature 2013, 500 (7463), 415–421.

8. Harris, R. S.; Dudley, J. P., APOBECs and virus restriction. Virology 2015, 479, 131–145.

9. Law, E. K.; Levin-Klein, R.; Jarvis, M. C.; Kim, H.; Argyris, P. P.; Carpenter, M. A.; Starrett, G. J.; Temiz, N. A.; Larson, L. K.; Durfee, C.; Burns, M. B.; Vogel, R. I.; Stavrou, S.; Aguilera, A. N.; Wagner, S.; Largaespada, D. A.; Starr, T. K.; Ross, S. R.; Harris, R. S., APOBEC3A catalyzes mutation and drives carcinogenesis in vivo. J Exp Med 2020, 217 (12), 20200261.

10. Sharma, S.; Patnaik, S. K.; Taggart, R. T.; Kannisto, E. D.; Enriquez, S. M.; Gollnick, P.; Baysal, B. E., APOBEC3A cytidine deaminase induces RNA editing in monocytes and macrophages. Nat Commun 2015, 6, 6881.

11. Maiti, A.; Hou, S.; Schiffer, C. A.; Matsuo, H., Interactions of APOBEC3s with DNA and RNA. Curr Opin Struct Biol 2021, 67, 195–204.

12. Petljak, M.; Dananberg, A.; Chu, K.; Bergstrom, E. N.; Striepen, J.; von Morgen, P.; Chen, Y.; Shah, H.; Sale, J. E.; Alexandrov, L. B.; Stratton, M. R.; Maciejowski, J., Mechanisms of APOBEC3 mutagenesis in human cancer cells. Nature 2022, 607 (7920), 799–807.

13. Jarvis, M. C.; Carpenter, M. A.; Temiz, N. A.; Brown, M. R.; Richards, K. A.; Argyris, P. P.; Brown, W. L.; Yee, D.; Harris, R. S., Mutational impact of APOBEC3B and APOBEC3A in a human cell line. bioRxiv 2022, 2022.04.26.489523.

14. Burns, M. B.; Lackey, L.; Carpenter, M. A.; Rathore, A.; Land, A. M.; Leonard, B.; Refsland, E. W.; Kotandeniya, D.; Tretyakova, N.; Nikas, J. B.; Yee, D.; Temiz, N. A.; Donohue, D. E.; McDougle, R. M.; Brown, W. L.; Law, E. K.; Harris, R. S., APOBEC3B is an enzymatic source of mutation in breast cancer. Nature 2013, 494 (7437), 366–370.

15. Harris, R. S., Cancer mutation signatures, DNA damage mechanisms, and potential clinical implications. Genome Medicine 2013, 5 (9), 1–3.

16. Leonard, B.; Hart, S. N.; Burns, M. B.; Carpenter, M. A.; Temiz, N. A.; Rathore, A.; Vogel, R. I.; Nikas, J. B.; Law, E. K.; Brown, W. L.; Li, Y.; Zhang, Y.; Maurer, M. J.; Oberg, A. L.; Cunningham, J. M.; Shridhar, V.; Bell, D. A.; April, C.; Bentley, D.; Bibikova, M.; Cheetham, R. K.; Fan, J. B.; Grocock, R.; Humphray, S.; Kingsbury, Z.; Peden, J.; Chien, J.; Swisher, E. M.; Hartmann, L. C.; Kalli, K. R.; Goode, E. L.; Sicotte, H.; Kaufmann, S. H.; Harris, R. S., APOBEC3B upregulation and genomic mutation patterns in serous ovarian carcinoma. Cancer Res 2013, 73 (24), 7222–7231.

17. Chan, K.; Roberts, S. A.; Klimczak, L. J.; Sterling, J. F.; Saini, N.; Malc, E. P.; Kim, J.; Kwiatkowski, D. J.; Fargo, D. C.; Mieczkowski, P. A.; Getz, G.; Gordenin, D. A., An APOBEC3A hypermutation signature is distinguishable from the signature of background mutagenesis by APOBEC3B in human cancers. Nat Genet 2015, 47 (9), 1067–1072.

18. Shi, K.; Carpenter, M. A.; Banerjee, S.; Shaban, N. M.; Kurahashi, K.; Salamango, D. J.; McCann, J. L.; Starrett, G. J.; Duffy, J. V.; Demir, Ö., Structural basis for targeted DNA cytosine deamination and mutagenesis by APOBEC3A and APOBEC3B. Nature Structural and Molecular Biology 2017, 24 (2), 131–139.

19. Kouno, T.; Silvas, T. V.; Hilbert, B. J.; Shandilya, S. M.; Bohn, M. F.; Kelch, B. A.; Royer, W. E.; Somasundaran, M.; Yilmaz, N. K.; Matsuo, H., Crystal structure of APOBEC3A bound to single-stranded DNA reveals structural basis for cytidine deamination and specificity. Nat Commun 2017, 8 (1), 1–8.

20. Silvas, T. V.; Schiffer, C. A., APOBEC3s: DNA-editing human cytidine deaminases. Protein Sci 2019, 28 (9), 1552–1566.

21. Hou, S.; Lee, J. M.; Myint, W.; Matsuo, H.; Kurt Yilmaz, N.; Schiffer, C. A., Structural basis of substrate specificity in human cytidine deaminase family APOBEC3s. J Biol Chem 2021, 297 (2), 100909.

22. Liu, M.; Mallinger, A.; Tortorici, M.; Newbatt, Y.; Richards, M.; Mirza, A.; van Montfort, R. L. M.; Burke, R.; Blagg, J.; Kaserer, T., Evaluation of APOBEC3B recognition motifs by NMR reveals preferred substrates. ACS chemical biology 2018, 13 (9), 2427–2432.

23. Harjes, S.; Jameson, G. B.; Filichev, V. V.; Edwards, P. J. B.; Harjes, E., NMR-based method of small changes reveals how DNA mutator APOBEC3A interacts with its single-stranded DNA substrate. Nucleic Acids Res 2017, 45 (9), 5602–5613.

24. Silvas, T. V.; Hou, S.; Myint, W.; Nalivaika, E.; Somasundaran, M.; Kelch, B. A.; Matsuo, H.; Kurt Yilmaz, N.; Schiffer, C. A., Substrate sequence selectivity of APOBEC3A implicates intra-DNA interactions. Scientific reports 2018, 8 (1), 7511.

25. Buisson, R.; Langenbucher, A.; Bowen, D.; Kwan, E. E.; Benes, C. H.; Zou, L.; Lawrence, M. S., Passenger hotspot mutations in cancer driven by APOBEC3A and mesoscale genomic features. Science 2019, 364 (6447), eaaw2872.

26. Brown, A. L.; Collins, C. D.; Thompson, S.; Coxon, M.; Mertz, T. M.; Roberts, S. A., Single-stranded DNA binding proteins influence APOBEC3A substrate preference. Scientific reports 2021, 11 (1), 21008.

27. Cortez, L. M.; Brown, A. L.; Dennis, M. A.; Collins, C. D.; Brown, A. J.; Mitchell, D.; Mertz, T. M.; Roberts, S. A., APOBEC3A is a prominent cytidine deaminase in breast cancer. PLoS Genetics 2019, 15 (12), e1008545.

28. Langenbucher, A.; Bowen, D.; Sakhtemani, R.; Bournique, E.; Wise, J. F.; Zou, L.; Bhagwat, A. S.; Buisson, R.; Lawrence, M. S., An extended APOBEC3A mutation signature in cancer. Nat Commun 2021, 12 (1), 1602.

29. Wörmann, S. M.; Zhang, A.; Thege, F. I.; Cowan, R. W.; Rupani, D. N.; Wang, R.; Manning, S. L.; Gates, C.; Wu, W.; Levin-Klein, R.; Rajapakshe, K. I.; Yu, M.; Multani, A. S.; Kang, Y.; Taniguchi, C. M.; Schlacher, K.; Bellin, M. D.; Katz, M. H. G.; Kim, M. P.; Fleming, J. B.; Gallinger, S.; Maddipati, R.; Harris, R. S.; Notta, F.; Ross, S. R.; Maitra, A.; Rhim, A. D., APOBEC3A drives deaminase domain-independent chromosomal instability to promote pancreatic cancer metastasis. Nat Cancer 2021, 2 (12), 1338–1356.

30. Barzak, F. M.; Harjes, S.; Kvach, M. V.; Kurup, H. M.; Jameson, G. B.; Filichev, V. V.; Harjes, E., Selective inhibition of APOBEC3 enzymes by single-stranded DNAs containing 2’-deoxyzebularine. Org Biomol Chem 2019, 17 (43), 9435–9441.

31. Kvach, M. V.; Barzak, F. M.; Harjes, S.; Schares, H. A.; Kurup, H. M.; Jones, K. F.; Sutton, L.; Donahue, J.; D’Aquila, R. T.; Jameson, G. B.; Harki, D. A.; Krause, K. L.; Harjes, E.; Filichev, V. V., Differential inhibition of APOBEC3 DNA-mutator isozymes by fluoro- and non-fluoro-substituted 2’-deoxyzebularine embedded in single-stranded DNA. ChemBioChem 2020, 21 (7), 1028–1035.

32. Kvach, M. V.; Barzak, F. M.; Harjes, S.; Schares, H. A. M.; Jameson, G. B.; Ayoub, A. M.; Moorthy, R.; Aihara, H.; Harris, R. S.; Filichev, V. V.; Harki, D. A.; Harjes, E., Inhibiting APOBEC3 activity with single-stranded DNA containing 2’-deoxyzebularine analogues. Biochemistry 2019, 58 (5), 391–400.

33. Kurup, H. M.; Kvach, M. V.; Harjes, S.; Barzak, F. M.; Jameson, G. B.; Harjes, E.; Filichev, V. V., Design, synthesis, and evaluation of a cross-linked oligonucleotide as the first nanomolar inhibitor of APOBEC3A. Biochemistry 2022, 61 (22), 2568–2578.

34. Serrano, J. C.; von Trentini, D.; Berríos, K. N.; Barka, A.; Dmochowski, I. J.; Kohli, R. M., Structure-guided design of a potent and specific inhibitor against the genomic mutator APOBEC3A. ACS chemical biology 2022, 17, 3379–3388.

35. Betts, L.; Xiang, S.; Short, S. A.; Wolfenden, R.; Carter, C. W., Jr., Cytidine deaminase. The 2.3 A crystal structure of an enzyme: transition-state analog complex. J Mol Biol 1994, 235 (2), 635–656.

36. Xiang, S.; Short, S. A.; Wolfenden, R.; Carter, C. W., Transition-state selectivity for a single hydroxyl group during catalysis by cytidine deaminase. Biochemistry 1995, 34 (14), 4516–4523.

37. Teh, A.-H.; Kimura, M.; Yamamoto, M.; Tanaka, N.; Yamaguchi, I.; Kumasaka, T., The 1.48 Å resolution crystal Structure of the homotetrameric cytidine deaminase from mouse. Biochemistry 2006, 45 (25), 7825–7833.

38. Xiao, X.; Li, S. X.; Yang, H.; Chen, X. S., Crystal structures of APOBEC3G N-domain alone and its complex with DNA. Nat Commun 2016, 7, 12193.

39. Maiti, A.; Myint, W.; Kanai, T.; Delviks-Frankenberry, K.; Rodriguez, C. S.; Pathak, V. K.; Schiffer, C. A.; Matsuo, H., Crystal structure of the catalytic domain of HIV-1 restriction factor APOBEC3G in complex with ssDNA. Nat Commun 2018, 9 (1), 2460.

40. Maiti, A.; Hedger, A. K.; Myint, W.; Balachandran, V.; Watts, J. K.; Schiffer, C. A.; Matsuo, H., Structure of the catalytically active APOBEC3G bound to a DNA oligonucleotide inhibitor reveals tetrahedral geometry of the transition state. Nat Commun 2022, 13 (1), 7117.

41. Grillo, M. J.; Jones, K. F. M.; Carpenter, M. A.; Harris, R. S.; Harki, D. A., The current toolbox for APOBEC drug discovery. Trends Pharmacol Sci 2022, 43 (5), 362–377.

42. Paar, M.; Schrabmair, W.; Mairold, M.; Oettl, K.; Reibnegger, G., Global regression using the explicit solution of Michaelis-Menten kinetics employing Lambert’s W function: high robustness of parameter estimates. ChemistrySelect 2019, 4 (6), 1903–1908.

43. Crooke, S. T.; Vickers, T. A.; Liang, X. H., Phosphorothioate modified oligonucleotide-protein interactions. Nucleic Acids Res 2020, 48 (10), 5235–5253.

44. Cummins, L. L.; Owens, S. R.; Risen, L. M.; Lesnik, E. A.; Freier, S. M.; McGee, D.; Guinosso, C. J.; Cook, P. D., Characterization of fully 2’-modified oligoribonucleotide hetero- and homoduplex hybridization and nuclease sensitivity. Nucleic Acids Res 1995, 23 (11), 2019–2024.

45. St. Martin, A.; Salamango, D. J.; Serebrenik, A. A.; Shaban, N. M.; Brown, W. L.; Harris, R. S., A panel of eGFP reporters for single base editing by APOBEC-Cas9 editosome complexes. Scientific reports 2019, 9 (1), 497.

46. McCann, J. L.; Salamango, D. J.; Law, E. K.; Brown, W. L.; Harris, R. S., MagnEdit— interacting factors that recruit DNA-editing enzymes to single base targets. Life Science Alliance 2020, 3 (4), e201900606.

## REFERENCES for METHODS

47. Research, G. Deprotection Supplement, Deprotection - Volumes 1-5. http://www.glenresearch.com/Technical/Deprotection.pdf.

48. Kabsch, W., Integration, scaling, space-group assignment and post-refinement. Acta Crystallogr D Biol Crystallogr 2010, 66 (Pt 2), 133–144.

49. Kabsch, W., XDS. Acta Crystallogr D Biol Crystallogr 2010, 66 (Pt 2), 125–132.

50. Vagin, A.; Teplyakov, A., Molecular replacement with MOLREP. Acta Crystallogr., Sect. D: Biol. Crystallogr. 2010, 66, 22–25.

51. Berman, H. M.; Westbrook, J.; Feng, Z.; Gilliland, G.; Bhat, T. N.; Weissig, H.; Shindyalov, I. N.; Bourne, P. E., The Protein Data Bank. Nucleic Acids Res. 2000, 28 (1), 235–242.

52. Murshudov, G. N.; Skubak, P.; Lebedev, A. A.; Pannu, N. S.; Steiner, R. A.; Nicholls, R. A.; Winn, M. D.; Long, F.; Vagin, A. A., REFMAC5 for the refinement of macromolecular crystal structures. Acta Crystallogr., Sect. D: Biol. Crystallogr. 2011, 67, 355–367.

53. Winn, M. D.; Ballard, C. C.; Cowtan, K. D.; Dodson, E. J.; Emsley, P.; Evans, P. R.; Keegan, R. M.; Krissinel, E. B.; Leslie, A. G. W.; McCoy, A.; McNicholas, S. J.; Murshudov, G. N.; Pannu, N. S.; Potterton, E. A.; Powell, H. R.; Read, R. J.; Vagin, A.; Wilson, K. S., Overview of the CCP4 suite and current developments. Acta Crystallogr., Sect. D: Biol. Crystallogr. 2011, 67, 235–242.

54. Emsley, P.; Lohkamp, B.; Scott, W. G.; Cowtan, K., Features and development of Coot. Acta Crystallogr D Biol Crystallogr 2010, 66 (Pt 4), 486–501.

55. Auerbach, A. A.; Becker, J. T.; Moraes, S. N.; Moghadasi, S. A.; Duda, J. M.; Salamango, D. J.; Harris, R. S., Ancestral APOBEC3B nuclear localization is maintained in humans and apes and altered in most other Old World primate species. mSphere 2022, 7, e00451–22.

