## Supplementary material for "Structure-guided inhibition of the cancer DNA-mutating enzyme APOBEC3A": Additonal analysis and discussion of results

Contents:

Supplementary Analysis and Discussion of Results

Figure S1

Tables S1-S2

### SUPPLEMENTARY ANALYSIS AND DISCUSSION OF RESULTS

Our structural studies establish that the stem-loop preconfigures the TC recognition motif in optimal position for binding to A3A, such that hairpin DNAs are more active substrates and as the 2′-deoxy-5-fluorozebularine derivative more potent inhibitor of A3A than linear ssDNA. Both 3- and 4-membered nucleotide loops present the TC motif in identical configuration. Although hinted at in earlier structures with ssDNA, a crucial role for His29 (and to a lesser extent Arg28) in substrate binding is confirmed, also explaining A3A preferential deamination of YTCR motif[16]. This has implications for activity of A3A leading to genomic instability and for the design of inhibitors to mitigate mutagenic activity of A3A in a wide variety of cancers. Interestingly, A3B does not have a strong preference in –2 position[17] and has an Arg instead of His29 in the corresponding position[16].

As the phosphorothioated derivatives, our inhibitors are highly resistant to nuclease degradation and can be directed to the nucleus with the aid of a commonly used transfection reagent. Moreover, we have obtained the first proof of inhibition of mutagenic activity of A3A in cells.

#### Isothermal Titration Calorimetric Results

Isothermal titration calorimetry (ITC) measurements were conducted for the binding of TTFdZ-hairpin (Extended Data **Fig. 4a**) and PS-TTFdZ-hairpin (Extended Data **Fig. 4b**) to A3A-E72A. Binding was substantially exothermic but offset by an unfavorable entropy of binding (Supplementary Information **Table 1**). At first sight this is counter-intuitive, as the hairpin is conformationally much less flexible than linear DNA; however, the negative entropy of binding reflects changes in dynamics of the protein and especially of the extended Loop-3 that in substrate-free structures is highly mobile. The dissociation constants $K_d$ cannot be compared with $K_m$ results, as the latter is on active enzyme that offers greater hydrogen-bonding potential than the former, as the former is on inactive enzyme A3A-E72A; additionally, to ameliorate problems of aggregation of protein due to the much higher concentrations required for ITC compared to enzymatic assays bovine serum albumin was present (30 mg/mL), as well as choline acetate (50 mM).

#### X-Ray Crystallographic Results

We obtained six structures, i.e. A3A-E72A with TTC-hairpin with and without zinc, ATTC-hairpin, CTTC-hairpin, and two structures of wild-type A3A with TTFdZ-hairpin. Notwithstanding full, partial or zero occupancy of the $Zn^{2+}$ site, all structures share a common binding of residues −2, −1, 0 and +1 to A3A-E72A, as highlighted in **Fig. 1e** and **f**. The nucleophilic water/hydroxide expected to be bound to the $Zn^{2+}$ center is replaced by $Cl^{-}$ in these structures and others of A3A-E72A, which all feature NaCl in the crystallization medium[19].

All structures are approximately isomorphous in space group $P2_1$. The packing of molecules in the unit cell is shown in cartoon form in Extended Data **Fig. 2**, highlighting differences in interactions between bases of hairpins for different hairpins All structures of A3A-E72A show a chloride in the active-site cavity, coordinating to the $Zn^{2+}$, when the latter is present. In addition to this chloride, there is a water molecule that hydrogen bonds to the chloride ion, to N3 of the cytosine at position 0, and to the main chain N of residue 72. This $Cl^{-}$ and $H_2O$ occupy, approximately, likely positions of the carboxylate oxygens of Glu72 of wild-type A3A (and also the structure of the catalytic C-terminal domain of A3G[20]). For structures of A3A, A3B and A3G where the general acid-base glutamate/glutamic acid has been mutated to alanine, this pair of atoms ($Cl^{-}$ and $H_2O$) is ubiquitous in electron density maps of all A3A, A3B and A3G enzymes, if not always included in the structural model structures of these enzymes[21]. For the structure of A3G with the Glu-->Ala mutation and substrate bound, a water molecule is reported at the site[22].

Consistent across all structures of A3 with single-stranded DNA oligonucleotide there is a tight turn to project substrate (or inhibitor) at position 0 into the active site. This is accomplished with non-standard torsion angles for the phosphate groups, such that phosphate...phosphate distances (as P…P) range from 6.2-7.3 Å for all nucleotides except for the phosphate group that links nucleotides at positions 0 and +1, where it shortens by more than an Ångstrom to 5.1-5.6 Å. The torsion angle for residues in the stem of the hairpin, $^{n}C3'$-$^{n}O3'$-$^{(n+1)}P$-$^{(n+1)}O5'$ and $^{n}O4'$-$^{n}C4'$-$^{n}C5'$-$^{n}O5'$, generally lie in the expected range for B-conformation DNA, respectively, ~-$90^{o}$ and $-70^{o}$·. Ultra-high resolution crystal structures (PDB ID 3u89 and 2be3)[23-24] of self-complementary 12-mer, ostensibly B-DNA, reveal considerable conformational flexibility and crystallographic disorder in the dodecamer with torsion angles well outside the canonical values for B-DNA. For residues in the loop (n = −2, −1, 0), the torsion angles $^{n}C3'$-$^{n}O3'$-$^{(n+1)}P$-$^{(n+1)}O5'$ are respectively approximately, $-160^{o}$, $+110^{o}$ and $60^{o}$, values substantially different to B-DNA.

For the phosphate linking positions -1 and 0, this amounts to an eclipsed conformation for the torsion angle C3¢-O3¢-P¢-O1¢. For residues 0 and +1 in three-membered loops, the torsion angles $^{n}$O4′-$^{n}$C4′-$^{n}$C5′-$^{n}$O5′ of ~176$^{o}$ also differ substantially from those generally expected for B-DNA and found in the stem of ~-70$^{o}$. Essentially the conformation changes from staggered to *trans*. For the four-membered loops the residue at −3 is flipped out and in order for the C-G pair at −4 and +1 to hydrogen bond, the torsion angle $^{n}$O4′-$^{n}$C4′-$^{n}$C5′-$^{n}$O5′ at residue +1 is ~+90$^{o}$. The torsion angle $^{n}$P-$^{n}$O5′-$^{n}$C5¢-$^{n}$C4¢ decreases from ~+170$^{o}$ in the stem and positions -2 and -1 to ~+120$^{o}$ at position 0.

All 2′-deoxyribose units adopt the expected C2′-endo conformation, with C5′-C4′-C3′-O3′ torsions angles in the range 120-150$^{o}$ (lower values generally associated with pyrimidine nucleobases and higher values with purine nucleobases, as previously observed[23-24]) except those at positions −4 and −3 in structures with four-membered loops, where to accommodate the flipped-out residue at -3, this torsion angle is ~85$^{o}$, a value characteristic of A-DNA.[25]

Our assignment of nitrogen atoms to the positions in the ring of His29 is based on the following interactions. His29-N$\delta_1$ (amine tautomer) forms a semi-salt bridge with the phosphate oxygen at the 3′ position of C$^{0}$, and also forms a sub-optimal hydrogen bond to deoxyribose O4′ of G$^{+1}$. An alternative assignment of orientation of His29 to form a canonical hydrogen bond between His29-N$\delta_1$ and O2 of T$^{-2}$ would require movement of His29 into a sterically less favourable conformation, as well as destroying the bifurcated hydrogen bond that locks in the sharp-turn. Moreover, the semi-salt bridge described above is much stronger than a canonical hydrogen bond between neutral parties in alternative assignment. In the highest resolution structure featuring the ATTC-hairpin, a well-defined water is observed off the assigned N$\varepsilon_2$ atom at 2.80 Å, which further confirms the assignment of atoms in His29.

*Inter alia*, O3¢ at position -1 forms a hydrogen bond to N$\delta_1$ of His29. His29-Nδ1 makes contact (3.4 Å) with ribose O4′ of guanine +1 and to phosphate O of cytosine 0, which in turns supports base pairing of G$^{+1}$ with C$^{-3}$. His29-Cδ2 makes a non-classical hydrogen bond with carbonyl oxygen O2 of thymine−2; and His29-Nε2 hydrogen bonds to a well-defined water. Finally, His29 pi stacks with G$^{+1}$, possibly as cation-π if His29 is protonated. Arg28 forms a cation-π interaction with T$^{-2}$.

Altogether, the A3A-hairpin interaction is provided by about 40 protein atoms and about 40 nucleotide atoms, highlighted as spheres with associated residues as sticks. Those are in van der Waals or hydrogen bonding contact at a tight threshold of 3.5 Å. A space-filling representation (Extended Data **Fig. 3a**) further highlights the key role of His29, Arg28 and Loop-3 in controlling the conformation of the loop and determining preference of A3A for purines in +1 and pyrimidines in –2 positions and for hairpin oligonucleotides over linear ssDNA.

#### Structure of A3A-E72A- ½$Zn^{2+}$ complexed with TTC-hairpin

This structure was determined to a resolution of 2.22 Å. The $Zn^{2+}$ ion is present in only about half occupancy. The chloride ion is present in full occupancy and is bound to the $Zn^{2+}$, if $Zn^{2+}$ is present. The $T^{-1}C^{0}N^{+1}$ (where N is G, T or A) moiety adopts a conformation very similar to that seen for linear oligo binding in one structure to A3A (PDB: 5keg[10]), but for binding in another A3A structure (5sww[21]) only the TC moiety closely aligns. All residues of the hairpin are observed, and the thymines at each foot of the stem hydrogen bond to each other (Extended Data **Fig. 2a,b**). In the previous structure of A3A-E72A (5keg) the cysteine 171 was mutated to alanine, most likely to prevent possible dimerization but in our structure Cys171 is present in reduced form. The interaction of TTC-hairpin with A3A-E72A is shown in Extended Data **Fig. 3c**.

#### Structure of A3A-E72A-no $Zn^{2+}$ complexed with TTC-hairpin

This structure was determined to a resolution of 2.10 Å. The $Zn^{2+}$ ion is entirely absent. Its absence is not an artefact of synchrotron radiation-induced damage, as an in-house data set from a crystal from the same drop also showed an absence of $Zn^{2}$. The chloride ion is present in full occupancy and located in the same position as in the $Zn^{2+}$-containing structures. The hairpin oligonucleotide superimposes closely on that for the (partially) $Zn^{2+}$-containing structure, A3A-E72A/TTC-hairpin. However, as illustrated in Extended Data **Fig. 1c**,**d**, there are substantial changes in positions of the otherwise $Zn^{2+}$-coordinating residues, as they seek to minimise repulsion. In particular, the loop bearing $Zn^{2+}$-binding Cys101 and Cyts106 flips to move these Cys away from each other and the other otherwise $Zn^{2+}$-binding ligand His70.

#### Structure of A3A-E72A-$Zn^{2+}$ complexed with ATTC-hairpin

The structure of A3A with a potentially 6-base-pair stem and a 4-membered loop was determined to 1.91-Å resolution. The T$^{-2}$TC motif binds the same as for the three-membered loop (**Fig. 1e**,**f**). However, the residue at −3 is flipped out, such that the cytosine at the −4 position hydrogen-bonds with little distortion to the guanine at +1 and the stem duplex is in register with substrates bearing just three residues in the loop (**Fig. 1e**,**f**). The chloride ion and water molecule noted earlier are also present in this structure. By virtue of being the structure with highest resolution, more water molecules are observed here than in other structures. The interaction of ATTC-hairpin with A3A-E72A is shown in Extended Data **Fig. 3** The packing of molecules is shown in Extended Data **Fig. 2d**,**e**. The low-resolution (3.15 Å) structure of A3A-E72A with CTTC-hairpin largely confirms this structure (**Fig. 1d**,**e**).

Whereas in structures with TTC-hairpin, which forms four canonical Watson-Crick base pairs and a thymine base pair, ATTC-hairpin has the potential to form an additional base-pair for a total of six Watson-Crick base pairs. However, five base pairs appear to define a “goldilocks” crystal packing. Thus, for ATTC-hairpin, in the crystalline state, there are only five base pairs and the last guanine ($^{+6}$G) interacts loosely with the $^{-8}$C$^{+5}$G pair of the second molecule in the asymmetric unit (Extended Data **Fig. 2d,e**). The cytosine at position −9 is not observed and is presumably flipped out and conformationally flexible.

### Structure of A3A-E72A-$Zn^{2+}$ complexed with CTTC-hairpin

Although this structure was determined to only 3.15 Å, the conformation of the oligonucleotide is well defined for residues −4 to +4. The shorter stem is less well defined, in part because four base pairs, one of which is an AT pair, do not confer high stability and in part because crystal packing is less effective where residues at the 5’ and 3’ ends of CTTC-hairpin base-stack weakly to the basal residues of the stem of the other A3A/CTTC-hairpin complex in the asymmetric unit, in contrast to the TTC- and ATTC-hairpins molecules (**Fig. 1d,e**). The 2′-deoxyribose-phosphate backbone of the loops is essentially identical to that for ATTC-hairpin, although the orientation of the flipped-out nucleobase at position −3 is possibly different. In both the ATTC- and CTTC-hairpin structures, this flipped-our nucleobase is very much more mobile than adjacent nucleobases, as evidenced by markedly higher atomic displacement parameters (*B*-values). In one subunit, the hairpin can be traced from −6 to +4 (end residue −7 not visible) and is superimposable on that for corresponding residues for ATTC-hairpin. However, in the other subunit, electron density for the hairpin is traceable from positions +6 to

−4 (relative to the expected conformation and threading of the hairpin analogous to that observed in the other subunit for all other hairpins). This leads to the uncomfortable conclusion that in order to maintain crystal packing contacts, this hairpin is threaded differently so that residues GAC**C**, rather than the expected CTT**C**, form the loop; that is, there is a three-residue shift. Relevant to this is that CCC, the preferred substrate of A3G, is a substrate of A3A, although a much poorer substrate than the preferred substrate TTC of A3A and A3B. Whereas in the expected threading a $C^{-4}G^{+1}$ pair sits at the top of the stem, in the unexpected threading it is a $C^{-7}T^{-2}$ pair (**Fig. 1d** numbering).

Amongst the various structures of A3A, there is always slight variations in the position of the B subunit (and its hairpin) relative to the A subunit, as seen in Extended Data **Fig. 1**.

#### Structure of wild-type A3A-Zn2+ complexed with TTFdZ-hairpin

This complex has been determined in two slightly different crystal forms of low isomorphism to 2.80 and 2.94 Å resolution (Extended Data **Table S2**). The molecules of the two forms and molecules A and B of the asymmetric unit of a given form are, however, closely isostructural (Extended Data **Fig. 1a**).

**Supplementary Table S1 | ITC data for titration of A3A-E72A with TTC- or PS-TTC-hairpin**[a]

| Experiment | n | K [$10^5/M$] | ΔH [*kcal/mol*] | ΔS [*cal/mol/K*] |
|---|---|---|---|---|
| PO-TTC-hairpin | $1.08 \pm 4.6 \times 10^{-3}$ | 8.72 ± 0.60 | -11.9 ± 0.073 | -12.6 |
| PO-TTC-hairpin | $1.03 \pm 2.9 \times 10^{-3}$ | 11.3 ± 0.56 | -12.6 ± 0.053 | -14.7 |
| PO-TTC-hairpin | $1.08 \pm 3.8 \times 10^{-3}$ | 6.72 ± 0.34 | -13.1 ± 0.066 | -17.3 |
| mean | 1.06 ± 0.024 | 8.91 ± 1.9 | -12.5 ± 0.49 | -14.9 ± 1.9 |
| PS-TTC-hairpin | $1.08 \pm 4.7 \times 10^{-3}$ | 6.62 ± 0.41 | -15.0 ± 0.092 | -23.8 |
| PS-TTC-hairpin | $1.04 \pm 4.1 \times 10^{-3}$ | 7.26 ± 0.42 | -15.6 ± 0.087 | -25.5 |
| PS-TTC-hairpin | $0.991 \pm 9.1 \times 10^{-3}$ | 3.87 ± 0.32 | -17.5 ± 0.22 | -33.2 |
| PS-TTC-hairpin | $1.03 \pm 6.8 \times 10^{-3}$ | 9.98 ± 0.10 | -14.9 ± 0.14 | -22.4 |
| mean | 1.03 ± 0.031 | 6.93 ± 2.2 | -15.8 ± 1.0 | -26.2 ± 4.2 |

[a] Error given is the fitting error except for the mean, where the root mean square is noted.

**Supplementary Table S2a | Kinetic constants determined for A3A-catalyzed deamination of linear and hairpin DNA, using NMR-based assay.**[a]

| Name | Sequence, 5'-to-3' | $k_{cat}$ (/s$^{-1}$) | $K_m$ (/μM) | $k_{cat}/K_m$ (/s$^{-1}$ mM$^{-1}$) |
|---|---|---|---|---|
| Linear DNA | $A_2T_2CA_4$ | 0.30 ± 0.06 | 3000 ± 900 | 0.10 ± 0.03 |
| TTC-hairpin | $T(GC)_2TTC(GC)_2T$ | 0.13 ± 0.03 | 31 ± 6 | 4.2 ± |

**Supplementary Table S2b | $K_i$ values of inhibitors of wild-type A3A against TTC-hairpin substrate.**[a]

| Name | Sequence, 5'-to-3' |
|---|---|
| FdZ-linear | 2400 ± 940 |
| TTFdZ-hairpin | 117 ± 15 |
| PS-TTFdZ-hairpin | 160 ± 70 |

[a] See Methods for experimental details.

**Supplementary Fig. S1**

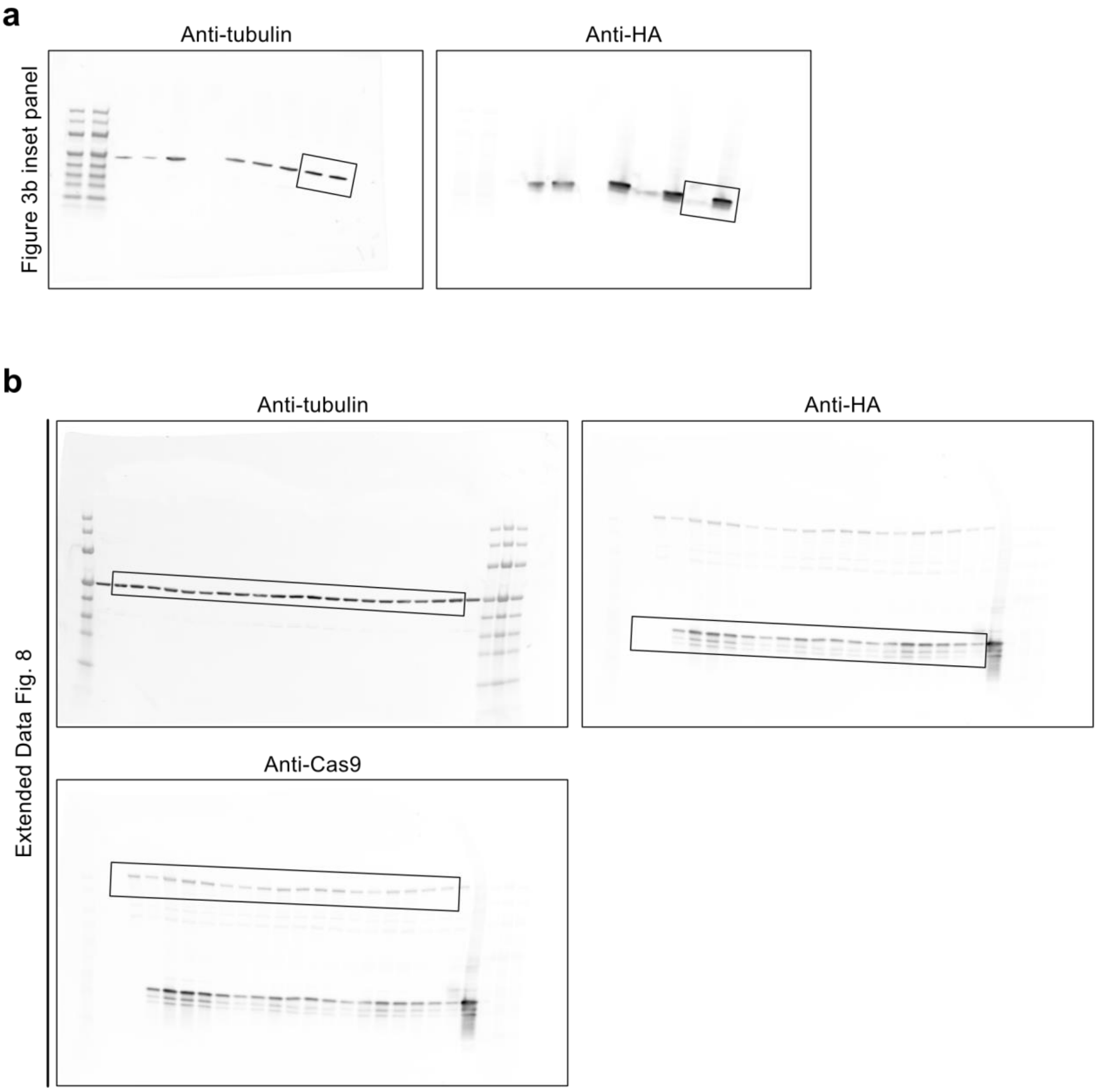


**Supplementary Fig. S1:** Original and uncropped immunoblots for (**a)** Fig. 3b inset panel and (**b**) Extended Data Fig. 8.
